# Fetal-like reversion in the regenerating intestine is regulated by mesenchymal Asporin

**DOI:** 10.1101/2021.06.24.449590

**Authors:** Sharif Iqbal, Simon Andersson, Ernesta Nestaite, Nalle Pentinmikko, Ashish Kumar, Daniel Borshagovski, Anna Webb, Tuure Saarinen, Anne Juuti, Alessandro Ori, Markku Varjosalo, Kirsi H. Pietiläinen, Kim B. Jensen, Menno Oudhoff, Pekka Katajisto

## Abstract

Epithelial tissues undergo fetal-like cellular reprogramming to regenerate after damage^1,2^. Although the mesenchyme and the extracellular matrix (ECM) play critical roles in tissue homeostasis and regeneration^2–5^, their role in repurposing developmental programs in epithelium is unknown. To model epithelial regeneration, we culture intestinal epithelium on decellularized small intestinal scaffold (iECM), and identify Asporin (Aspn), an ECM bound proteoglycan, as a critical mediator of cellular reprogramming. Aspn is produced by the mesenchyme, and we show that its effect on epithelial Tgfβ-signalling via CD44 is critical for fetal-like conversion. Furthermore, we demonstrate that *Aspn* is transiently increased upon chemotherapy-induced damage and pivotal for a timely induction of the fetal-like state and tissue regeneration. In summary, we establish a platform for modelling epithelial injury responses *ex vivo*, and show that the mesenchymal *Aspn*-producing niche controls tissue repair by regulating epithelial fetal-like reprogramming.

---

Cellular plasticity is integral for intestinal regeneration as a mechanism for restoring epithelial integrity following damage such as ulceration and mucositis^2,6^. Intestinal organoids^7^ provide a useful tool to study the intra-epithelial mechanisms of this process^8^, however, organoids are typically grown in reconstituted matrices that are distinct from the native environment of the epithelial cells. Moreover, *in vivo* repair involves many extracellular matrix components^2^ and inputs from the mesenchyme^9–11^. Hydrogel matrices that support organoid culture lack the native ECM composition and factors produced by the stroma^12^ and do not allow studies probing on their function during regeneration. We reasoned that tissue decellularization, which removes live cells while retaining matrix bound growth factors and the acellular ECM architecture intact, may provide an attractive complement to the organoid system as a native scaffold that includes many of the cues guiding regeneration *in vivo*. Unlike the synthetic scaffolds^13^, decellularized intestine contains the organization of the basement membrane and auxiliary ECM factors, and akin to tissue regeneration following damage provide a surface substrate for tissue remodelling independent of dynamic cellular inputs from the stromal compartment.

In order to develop a model that allows quantification of the epithelial reparative growth and the impact of extra-epithelial mechanisms, we developed a functional assay using decellularized small intestinal matrixes (iECM). We expanded on published decellularization protocols^12^ to allow efficient growth and expansion of epithelial cells *ex vivo* **(Extended Data Fig. 1a)**. Decellularization preserved the 3D tissue architecture with distinct regions of former villi and crypts **(Fig.1a, Extended Data Fig. 1b)**. When we re-introduced epithelium onto the iECMs by seeding either freshly isolated mouse small intestinal crypts or organoids^7^, the seeded epithelial cells grew from small islands to form a monolayer covering both the empty crypt pits and denuded villi with striking similarity to the organization and topology of the live tissue (**Fig.1a-c, Extended Data Fig. 1b**). As in the native tissues, Keratin 20, a marker of villous differentiation^14^ was restricted to the epithelium covering villi (**Fig.1b**), and proliferating cells marked by EdU incorporation were confined to the re-epithelialized crypts (**Ext. Data Fig. 1c**). Furthermore, when cells from the *Lgr5-EGFP-IRES-CreERt2* reporter mouse was used for the re-epithelialization of the iECMs Lgr5+ cells, and Lyz+ Paneth cells were restricted at the crypt bottom, as in the native epithelium (**Fig. 1b & Ext. Data Fig. 1d**). Furthermore, the fully re-epithelialized iECMs reached a steady state, where stem and progenitor cells proliferate in steady crypts, – and differentiating cells move to villi. This allowed prolonged culture (>3 weeks), as the dead cells shedding from villus can be removed from the open layout iECMs by media change. This is in contrast with intestinal organoids, where differentiated epithelial cells exfoliate into the organoid lumen, and it is therefore necessary to passage crypt domains of mechanically disrupted organoids to ensure continuous propagation. Demonstrating self-renewal capacity, crypts isolated from the re-epithelialized iECM formed organoids in Matrigel similarly to crypts isolated directly from the mouse intestine **(Ext. Data Fig. 1e)**. Jointly, these findings reveal that signals associated with the 3D architecture of the tissue including extracellular matrix components guide cell fate, and that stem cells self-renew on the iECM allowing analysis of re-epithelization of exposed ECM *ex vivo.*

**Figure 1.**
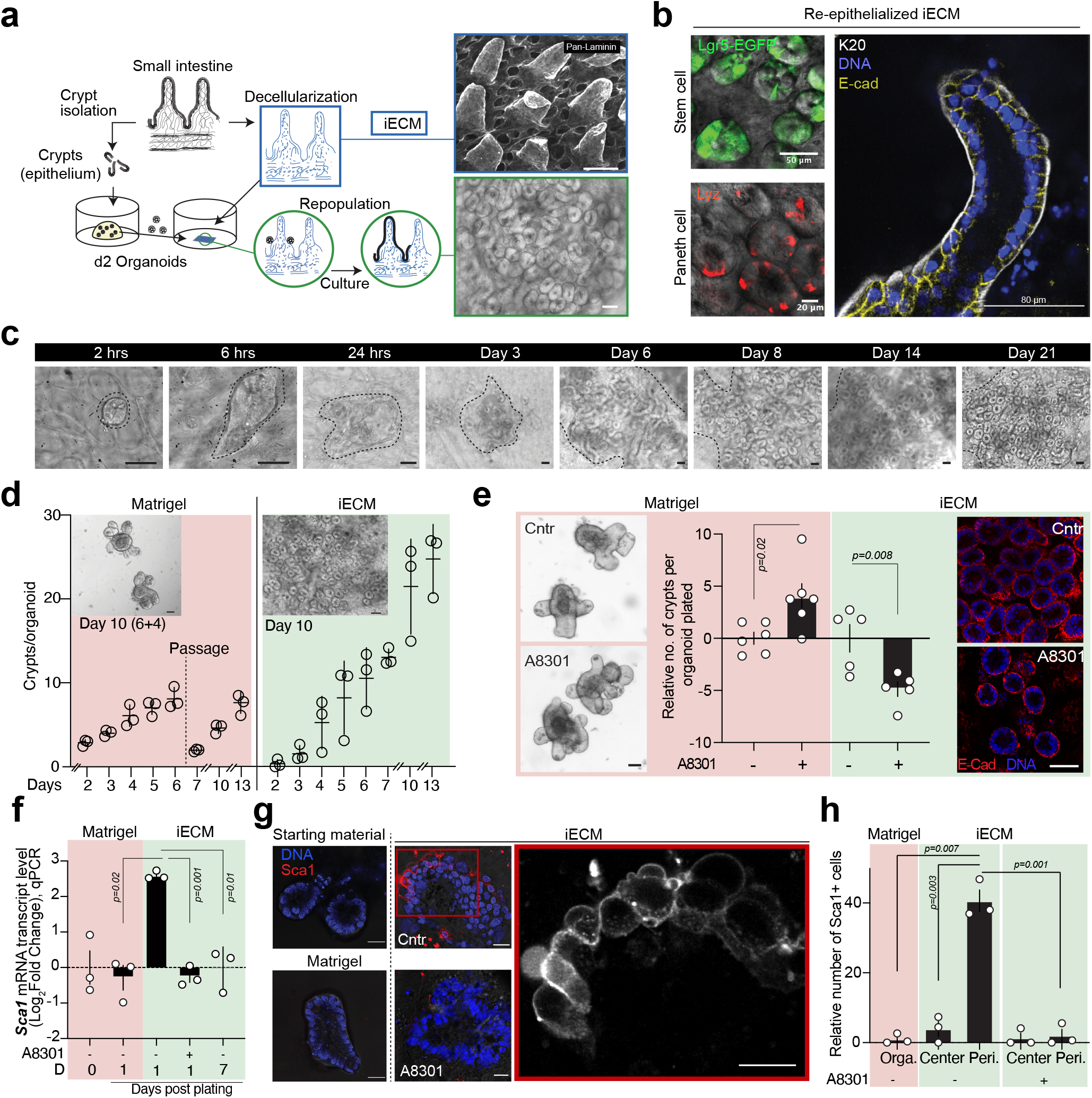
Small intestinal epithelium adopts Tgfβ-dependent fetal-like pro-regenerative program to regenerate on decellularized ECM (iECM). **a**, Generation of decellularized intestinal ECM scaffolds (iECM). Immunofluorescense image of the Pan-Laminin stained iECM. Scale bar 100 μm (top; iECM) and 50 μm (down; repopulated iECM). **b**, Immunofluorescencent detection of differentiated villous epithelium (Keratin20, K20; E-cadherin, E-cad), stem cells (GFP) and Paneth cells (Lysozyme, Lyz) on iECM repopulated with *Lgr5-CreERt2-IRES-EGFP* epithelium for 8 days. **c**, Culture of the intestinal crypts on iECM from the single intestinal organoid on iECM at different timepoints. Scale bar 50 μm. **d**, Comparison of the growth dynamics of intestinal epithelium in organoid culture and on iECM. Organoid culture (red); iECM (green). n=3 mice. mean +/− s.d. Scale bar 50 μm. **e**, Impact of the inhibition of Tgfβ receptor Type I (TgfβRI) on the regenerative capacity of the intestinal epithelium in matrigel and on iECM. Representative micrographs show the growth of new crypts from single spherical organoid in five (matrigel; left) and seven (iECM; right) days. E-cad (red); DNA (blue). A8301 = TgfβRI inhibitor. Student’s paired t-test, mean +/− s.d. Scale bar 50 μm. **f**, qPCR analysis of the relative mRNA level of *Sca1* from the intestinal organoids (D0: before passaging; D1: 24 hrs after passaging in matrigel) and intestinal epithelium on iECM (D1 & D7: Day 1 & Day 7 post-plating on iECM from matrigel). *Rpl13a* was used as a reference gene. Values show fold change in comparison to Day 0 intestinal organoids. Student’s paired t-test, mean +/− s.e.m. **g**, Representative immunofluorescent images of the intestinal organoids, and intestinal epithelium (Day 1 and Day 7 post-plating) on iECM. Scale bar 20 μm. **h**, Relative number of Sca1+ cells in the matrigel-cultured intestinal organoids, and in the intestinal epithelium plated on iECM (day 1 post-plating). Values were normalized with the number of Sca1+ cells in the intestinal organoids. Student’s paired t-test, mean +/− s.e.m. Peri. = Peripheral epithelial cells on iECM. n= 3 mice.

Even though the growth rate of epithelial cells on iECM was comparable to organoids in Matrigel (**Fig. 1d**), the process of iECM re-epithelialization was distinct from the growth of organoids. Whereas new crypts in organoids appear by self-organization of the epithelium^7^, the epithelial monolayer growing on iECM formed crypts only into the predetermined empty crypt pits of the former tissue. As the epithelial growth on iECM was guided by its intact ECM, we asked whether re-epithelialization of the empty iECM *ex vivo* recapitulates aspects of *in vivo* re-epithelization following damage. Epithelial cells participating in tissue regeneration transition into a fetal-like state expressing markers including *Sca1^2^* and *Clu^15^*. Interestingly, epithelial cells seeded on iECM displayed a profound upregulation of both markers (**Fig. 1f; Ext. Data Fig. 2b**), suggesting that the seeding on iECM recapitulates the epithelial response during tissue regeneration.

Damage-induced developmental reversion in the epithelium can be triggered Yap/Taz dependently by exposure to ECM constituents^2^, and by interferon gamma in response to parasitic infection^1^. Tgfβ signalling is another pathway critical for epithelial homeostasis^16^ and regeneration^11,17,18^, and has been linked to developmental reversion in cancer cells^19^. In order to probe the importance of Tgfβ signalling in re-epithelization of the iECM and in the conversion from a homeostatic to a regenerative phenotype, we used the Tgfβ Type I receptor (TgfβRI) kinase inhibitor A8301^20^. In Matrigel embedded organoids A8301 increased the crypt formation significantly, whereas on iECM A8301 reduced the re-epithelization and *de novo* crypt formation (**Fig.1e**). As A8301 induced opposite effects in epithelium cultured in Matrigel and on iECM, we next assayed the Tgfβ signalling activity on the two culture systems. Expression of target genes of Tgfβ-signalling was increased when dissociated crypt domains from Matrigel grown organoids were placed on the iECM (**Ext. Data Fig. 2a**). Importantly, the significant burst of fetal-state markers *Sca1* and *Clu* on iECM was completely dependent on Tgfβ signalling (**Fig.1f; Ext. Data Fig. 2b**). Moreover, the transient induction of the Sca1+ fetal-like cells was restricted to the periphery of the expanding epithelium (**Fig.1g-h**). Jointly these data suggest that a Tgfβ induced fetal program is integral to the re-epithelization of the decellularized iECM scaffolds.

To identify which Tgfβ pathway modulating factors are retained in the iECM, and could contribute to the Tgfβ dependent engagement of the epithelial fetal-like program, we examined the proteomic composition of iECMs by mass spectrometry. Indicating that the decellularization process effectively removes majority of the intracellular proteins, 15 of the 20 most abundant proteins were associated with the ECM (matrisome database^21^) **(Table-1)**. Among these, we identified Decorin and Asporin, two Tgfβ pathway modulating short leucine rich repeat proteoglycans (SLRPs)^22^, and another Tgfβ pathway modulating SLRP, Biglycan, was also detected with lower abundancy. Neither recombinant Decorin (1 μg/ml) or Biglycan (1 μg/ml) induced Tgfβ -signaling and fetal-like markers when tested on intestinal organoids (**Ext. Data Fig. 3).** In contrast, purified recombinant asporin (Aspn) (**Ext. Data Fig. 4a**) strongly stimulated expression of Tgfβ-responsive genes in intestinal organoids (**Fig. 2a, b**). *In situ* hybridization analysis of intact tissues revealed that *Aspn* is detected in pericryptal mesenchymal cells (**Fig. 2c**), which have been shown to support epithelial functions in homeostasis and tumorigenesis by secreting factors modulating Tgfβ, Bmp, Yap and Wnt signalling in the epithelium^3,4,23^. Consistently, by re-analysing the publicly available data, we observed high level of *Aspn* expression in *Pdgfrα^low^* population which reside around the crypt base^4^ (**Ext. Data Fig. 4b**).

**Figure 2.**
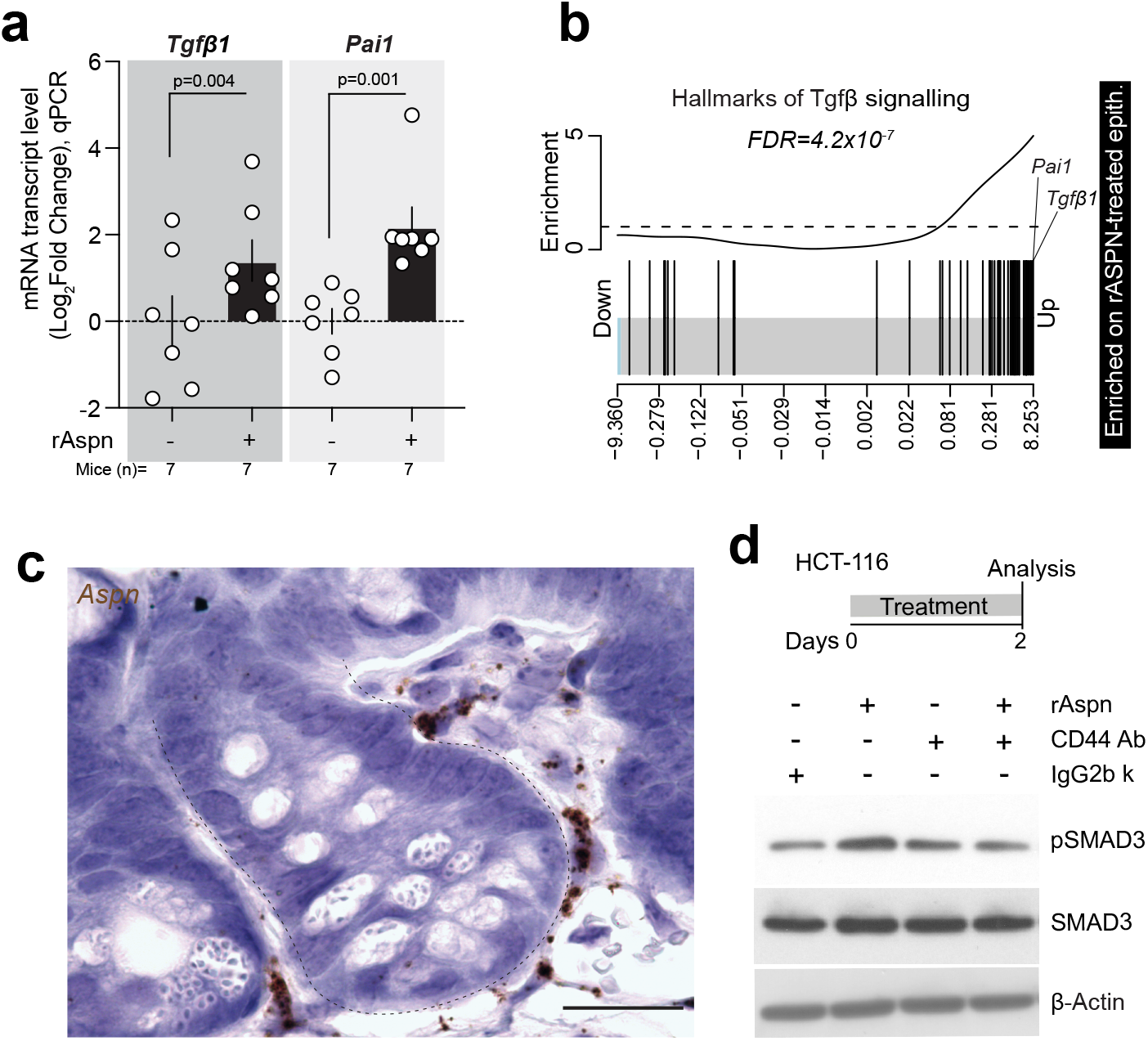
Mesenchymal Aspn promotes Tgfβ signaling via CD44 receptor. **a**, qPCR analysis of *Tgfβ1 and Pai1* expression in the rAspn-treated (48 hrs) intestinal organoids. Values show fold change in comparison to control. *Rpl13a* was used as a reference gene. Student’s paired *t*-test, mean +/− s.e.m. **b**, GSEA analysis for the gene list “Hallmarks of Tgfβ signaling” on transcription profiles from rAspn (500ng/ml) treated (48 hrs) organoids. False discovery rate (FDR) of enrichment is shown. **c**, *In situ* analysis of *Aspn* expression in the pericryptal area. Brown (Aspn). Scale bar 20 μm. **d**, Immunoblot for pSMAD3, SMAD3 and β-ACTIN on rAspn and/or CD44 Ab-treated (48 hrs) HCT-116 cells.

Extracellular Aspn is reported to bind Tgfβ ligands and thereby inhibit downstream Tgfβ signalling^24^. However, we noted that rAspn promoted Tgfβ signalling. Therefore, we investigated other possible mechanisms modulating the Aspn-Tgfβ signalling axis. Aspn binds the CD44 transmembrane hyaluronan receptor^25,26^, which is highly expressed by the intestinal stem cells and progenitors^27^ and increased in the fetal-like regenerative population^1^. Via CD44, Aspn can activate epithelial mesenchymal transition (EMT) and NF-κB pathways^26^, but interestingly, CD44 can also physically interact with Tgfβ receptor I and activate Tgfβ/Smad signalling Tgfβ-ligand independently^28^. Consistently, we observed that rAspn treatment of the intestinal organoids led to significant changes in genes responsive to CD44-downstream pathways including Tgfβ **(Fig 2b)**, EMT, Stat3 and NF-κB^28,29^ **(Ext. Data Fig. 5a-c**). rAspn treatment also modestly increased the Yap/Taz pathway regulated genes (**Ext. Data Fig 6**). These findings suggested that Aspn promotes Tgfβ signalling in the intestine via the CD44 transmembrane receptor. In support, CD44 function blocking antibody^30^ blunted the effects of rAspn on Tgfβ signalling **(Fig. 2d).**

Since Aspn was abundant in the iECM, and rAspn induced Tgfβ signalling, we next asked whether the Aspn-induced Tgfβ signalling underlies the epithelial fetal-like reversion that we observed on the iECM during tissue reepithelization. We first tested whether rAspn can induce the fetal-state in the intestinal organoids in Matrigel. Excitingly, rAspn reverted a significant portion of the intestinal organoids into Sca1+ spheroids similar to what has been observed from fetal intestinal epithelium (**Fig. 3a**). Moreover, the transcriptional changes induced by rAspn overlapped strikingly with the previously reported transcriptional profile of fetal intestinal organoids^31^ **(Fig. 3b)**.

**Figure 3.**
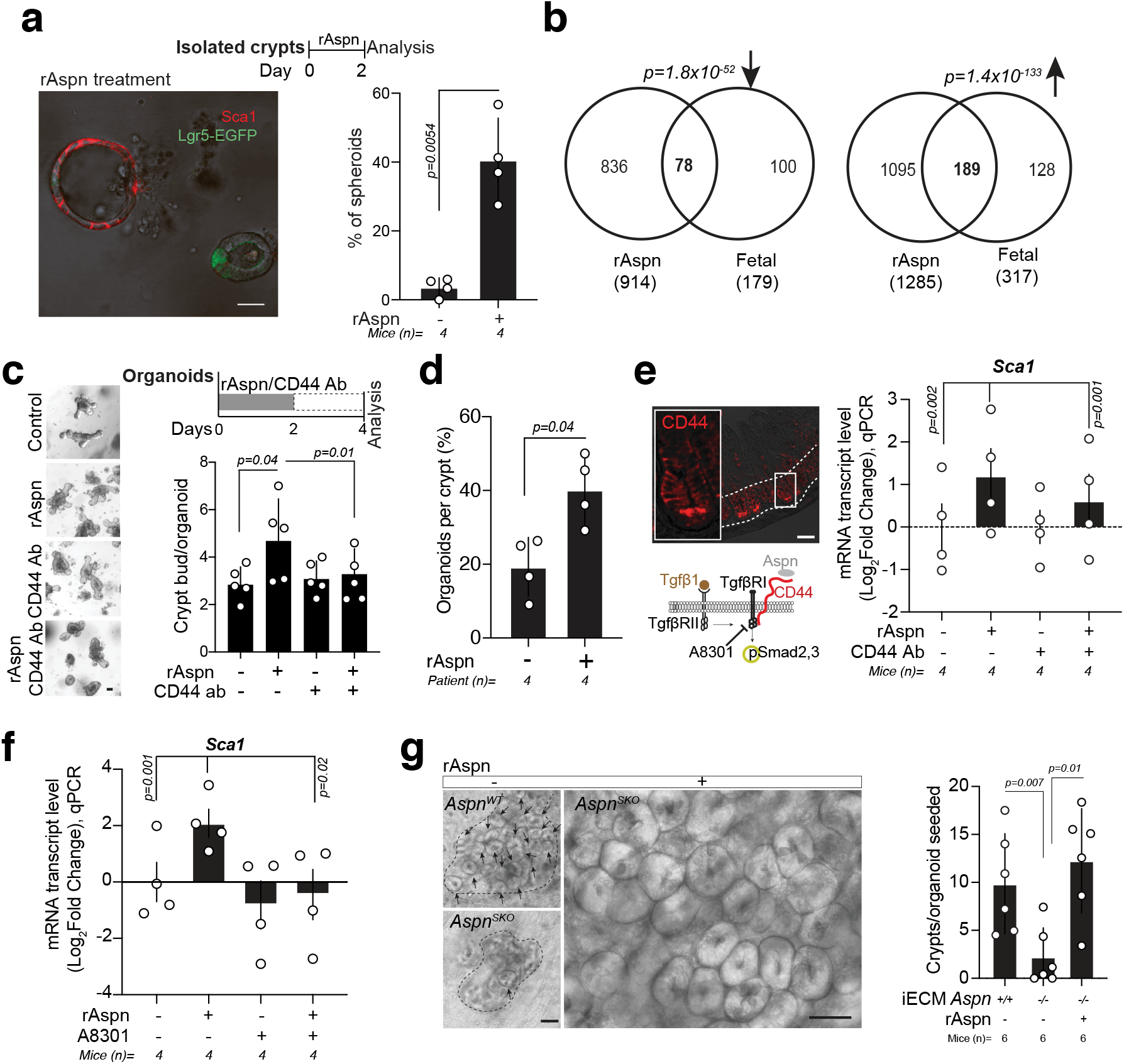
rAspn promotes intestinal epithelial regeneration by inducing fetal-like transcriptional state via Tgfβ signalling. **a**, Analysis of spheroidicity of rAspn-treated organoids (n=4). Crypts were isolated from *Lgr5-CreERt2-IRES-EGFP* reporter mice. Mean +/− s.d. Student’s paired t-test. Lgr5+ (Green) stem cells and Sca1 (Red) staining are shown in the rAspn-treated organoids (left). Scale bar 30 μm. **b**, Venn diagram of genes with altered expression in mouse fetal organoids and in adult organoids after rAspn treatment. *P*-values show significance of overlap. Upregulated (Upward arrow), downregulated (Downward arrow). **c**, Regenerative growth of crypts (n=5 mice) treated transiently with rAspn, and/or CD44 function-blocking antibody/IgG2b kappa control antibody. Mean +/− s.d. Student’s paired t-test. Scale bar 50 μm. **d**, Organoid forming capacity of rAspn treated isolated human small intestinal crypts (n=4 human subjects). Mean +/− s.d. Student’s paired t-test. **e-f**, qPCR analysis of the relative mRNA level of *Sca1* in the rAspn treated mouse intestinal organoids (n=4 mice). CD44 function-blocking antibody or isotype control IgG2b kappa (in **e**), and Tgfβ Type I receptor inhibitor (A8301) (in **f**), were used to probe dependency on CD44 and TgfβRI. Sites of intervention and CD44 staining in the intestinal epithelium are shown in **e**. Values show the fold change in comparison to untreated control organoids. *Rpl13a* was used as a reference gene. Mean +/− s.e.m. Student’s paired *t*-test. **g**, Regenerative growth of the wild type epithelium on *Aspn^WT^* and *Aspn^SKO^* iECM with or without rAspn (500 ng/ml; 2 days treatment). Mean +/− s.d. Student’s paired *t*-test. Scale bar 50 μm.

We tested if the fetal-like spheroidal growth induced by Aspn can promote the formation new crypts in organoids. After withdrawal of rAspn, fetal-like spheroids were resolved into budding organoids, and transient pulse of rAspn increased crypts formation in organoids in a dose-dependent manner **(Ext. Data Fig. 7)**. Furthermore, compared to the IgG2b κ treated control organoids, the effect of rAspn on crypt formation in organoid was blunted by the CD44 blocking antibody **(Fig. 3c)**. Transient rAspn also increased the organoid forming capacity of human small intestinal crypts, demonstrating that effects of Aspn on intestinal epithelium are conserved **(Fig. 3d)**. However, while CD44-blocking antibody reduced the transcriptional effects of rAspn on fetal markers significantly **(Fig 3e; Ext. Data Fig. 8a)**, only A8301 fully blocked the effects of Aspn (**Fig. 3f; Ext. Data Fig. 8b-d**). This suggested that CD44 blockade with an antibody is not complete, and may initiate the positive feedback-loop activating the Tgfβ-pathway via *Tgfβ1* expression^32^. We therefore tested whether Tgfβ1 can directly induce fetal-like state in organoids. Recombinant Tgfβ1 (0.1 ng/ml) induced fetal-like spheroid formation in organoids similarly to Aspn (**Ext. Data Fig. 9a-c)**, increased crypt formation in organoids when administered transiently (48h), but decreased it upon sustained treatment **(Ext. Data Fig. 9d).** Taken together, Aspn induces fetal-like regenerative state in the epithelium via CD44 and the Tgfβ-pathway, and its effects can be recapitulated by Tgfβ-ligand mediated pathway activation.

Cross-talk between mesenchyme and epithelium is necessary for *in vivo* homeostasis ^3,4,33^ and tissue repair ^9–11^. To study the role of mesenchymal Aspn in epithelial regeneration on iECM *ex vivo*, and *in vivo*, we generated a mouse model harbouring a conditional allele of *Aspn* (*Aspn^lox^*) **(Ext. Data Fig. 10a)**, and crossed it with *Twist2-Cre* mice^34^ to delete *Aspn* in the mesenchyme (*Twist2-Cre; Aspn^lox/lox^*, hereafter *Aspn^SKO^*) (**Ext. Data Fig. 10a-f**). *Aspn^SKO^* mice are fertile and develop normally, allowing us to generate comparable iECMs from *Aspn^WT^* and *Aspn^SKO^* mice. In comparison to *Aspn^WT^* iECM, wild type epithelium grew significantly slower on *Aspn^SKO^* iECM (**Fig. 3g**). However, epithelial regeneration on the *Aspn^SKO^* iECM was rescued with transient exogenous rAspn treatment in the start of the culture (**Fig. 3g**). These data demonstrate that stromally deposited Aspn supports regeneration on iECM.

We further probed the role of Aspn during tissue repair *in vivo* by administering mice with 5-Fluorouracil (5-FU). In mice, acute 5-FU treatment induces cell death in crypts leading to loss of body weight due to reduced water retention and nutrient intake^35^, and provides a tractable system for assessing intestinal injury, repair, and recovery^36^. We observed that *Aspn* expression is transiently increased after 5-FU (200 mg/kg) **(Fig. 4a)**, but subsides during the later stages of intestinal regeneration (day 5 post 5FU). Furthermore, when treated with 5-FU, the *Aspn^SKO^* mice did not recover like WT animals **(Fig. 4b)**. Importantly, the initial weight loss was similar to *Aspn^WT^* mice, suggesting that *Aspn^SKO^* mice experience similar damage as *Aspn^WT^* mice, but that tissue repair and regeneration is impaired. Consistent with the notion of poor regeneration, cellular density of the villi **(Fig. 4c)**, and villus/crypt -length ratio **(Ext. Data Fig. 11)** were reduced in *Aspn^SKO^* at five days after 5-FU.

**Figure 4.**
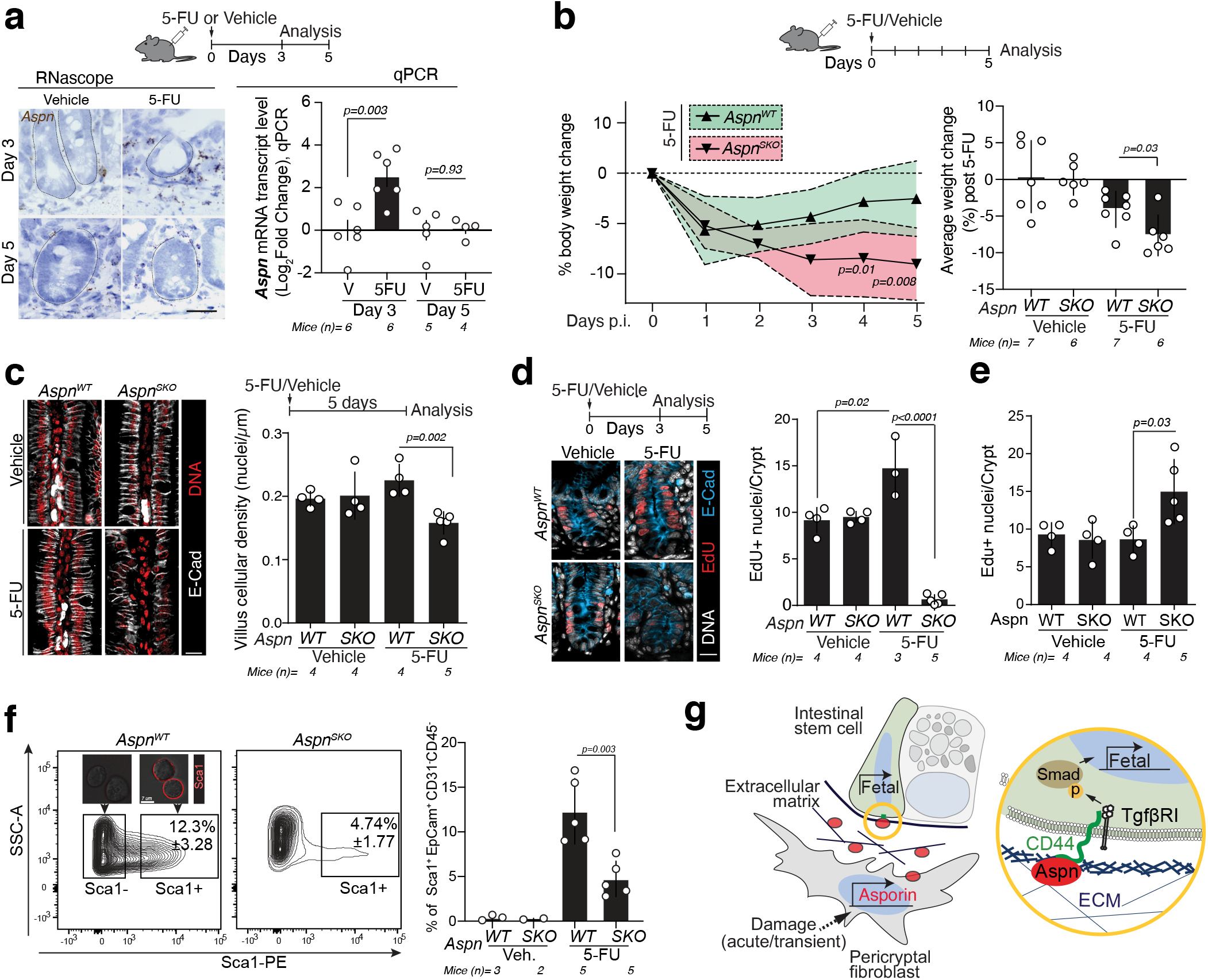
Mesenchymal loss of Aspn impedes induction of epithelial fetal-like state and repair after damage. **a**, Analysis of Aspn expression in the mouse intestine three and five days after 5-Fluorouracil injection (5-FU, 200 mg/kg). *In situ* hybridization (left) and qPCR (right). qPCR values show the fold change relative to Vehicle (V; DMSO)-treated mice. *Actin* as a reference gene in qPCR. Scale bar 20 μm. Student’s unpaired *t*-test. **b**, Relative body weight loss of *Aspn^WT^* and *Aspn^SKO^* mice (n=6-7 mice per group) treated with vehicle (DMSO) or 5-FU. Daily data points represent median and interquartile range (dashed line). Average weight loss post 5-FU (days 1-5) is shown for all groups (bar graph, right). Student’s unpaired *t*-test. **c**, Cellular density of the ileal villi in *Aspn^WT^* and *Aspn^SKO^* mice after vehicle or 5-FU (n=4-5 mice/group). Scale bar 20 μm. **d-e**, Quantification of EdU+ proliferative epithelial cells in *Aspn^WT^* and *Aspn^SKO^* mice three (**d**) and five (**e**) after vehicle/5-FU injections (n=3-5 mice per group analysed). Scale bar 20 μm. Mean +/− s.d. Student’s unpaired *t*-test. **f**, Flow cytometry analysis of small intestinal Sca1+EpCam+CD31^−^CD45^−^ cells isolated from vehicle and 5-FU treated *Aspn^WT^* and *Aspn^SKO^* mice (n=2-5; Day 3 post injection). FACS-sorted Sca1-EpCam+CD31^−^CD45^−^ (inset; left) Sca1+EpCam+CD31^−^CD45^−^(inset; right), and average frequency (+/− s.d.) of Sca1+EpCam+CD31^−^CD45^−^ cells is shown (n=2-5 mice). Bar graph shows the frequency of Sca1-Ep-Cam+CD31^−^CD45^−^ cells (n=2-5). Mean +/− s.d. Student’s unpaired *t*-test. **g**, Schematic model on the role of Aspn in intestinal regeneration.

To further address why *Aspn^SKO^* mice recover poorly, we assessed proliferation in the 5-FU damaged epithelium. Whereas in *Aspn^WT^* mice proliferation peaks at day 3 post injection and recedes back to normal level at day 5 post injection **(Ext. Data Fig. 12)**, *Aspn^SKO^* mice contrasted this pattern with dramatic reduction in proliferation at 3 days **(Fig. 4d)** and increased proliferation at five days after 5FU **(Fig. 4e)**. Jointly these data suggest that loss of stromal *Aspn* retards the engagement of the regenerative program in the epithelium.

Finally, we sought to investigate Tgfβ pathway activity and cell state reversion in *Aspn^WT^* and *Aspn^SKO^* mice during 5-FU induced regeneration. In line with normal development and intestinal histology of the unchallenged *Aspn^SKO^* mice, pSmad2 levels in the crypt epithelium of the vehicle treated *Aspn^SKO^* and *Aspn^WT^* mice were similar **(Ext. Data Fig. 13)**. However, crypts in *Aspn^WT^* mice had significantly higher peak pSmad2 levels upon damage (day 3 post 5-FU) that subsided to normal levels upon recovery (day 5 post 5-FU) **(Ext. Data Fig. 13)**. The reduced initial induction of Tgfβ signalling in *Aspn^SKO^* mice also resulted in longer maintenance of damage-induced Tgfβ activity, supporting the notion of late onset of proliferation **(Fig. 4d-e)** and delayed tissue repair. These data confirm that injury-induced mesenchymal Aspn can promote rapid activation of epithelial Tgfβ signalling *in vivo*, and the Aspn-mediated temporally controlled boost to Tgfβ induction is necessary for proper epithelial regeneration. Importantly, frequency of Sca1^+^ cells was dramatically reduced in *Aspn^SKO^* mice three days after 5-FU **(Fig. 4f; Ext. Data Fig. 14)**, suggesting that defects in the CD44-TgfβRI mediated induction of the fetal-like regenerative state delays epithelial regeneration after deletion of mesenchymal *Aspn*.

The culture platform we describe here allows *ex vivo* recapitulation of early epithelial injury responses and overcomes many shortcomings of closed-format organoids. Using this platform, we discovered a novel mesenchymally produced intestinal niche factor Asporin, which upon tissue damage induces a transient fetal-like change in the epithelial cell state and thereby coordinates tissue repair. Taken together, this study highlights the dynamic nature and importance of cross-talk between mesenchymal and epithelial cells during regeneration, and suggests that the Aspn-CD44-Tgfβ signalling axis has evolved to allow temporal control over tissues repair programs. The Aspn-mediated regulation of epithelial cell-state may also provide new opportunities to target epithelial tumor initiation^23^ and ulcerative colitis^2^, where drift of developmental programs is implicated.

## METHODS

### Isolation of mouse intestinal crypts

Mouse small intestinal crypts were isolated as previously published^37^. Briefly, mouse small intestine was flushed with the cold PBS, mucus and mesentery were removed, and subsequently, cut opened. Intestine was cut into smaller pieces and incubated with 10 mM EDTA in PBS (3x) on ice for 2 hr. Epithelium was detached by vigorous shaking, and filtered through 70 μm nylon mesh. Enriched crypts were washed with cold PBS once more, and plated in 60% Matrigel (BD Biosciences) with ENR media. 10 μM Y-27632 was added to the media for the first 2 days.

### Isolation of human crypts

Human small intestinal biopsies were cut into small pieces on ice cold PBS. Biopsies are then incubated with 10 mM EDTA in PBS (with 3x changes) on ice for 2 hr. Crypts were isolated by vigorously shaking, and filtered through 70 μm nylon mesh. Isolated crypts were washed with ice cold PBS and cultured in 60% Matrigel (BD Biosciences) as described previously^38^.

### Organoid culture

200-300 crypts were plated per 20μl drop of 60% Matrigel and overlaid with ENR media (DMEM/F12 (Gibco), 1x Glutamax (Gibco), 100 U/ml of Penicillin and Streptomycin, 10 mM Hepes, 50 ng/ml of mouse EGF (RnD), 100 ng/ml noggin (Peprotech), 500 ng/ml of RSpondin-1 (RnD), 1 μM N-Acetyl-L-cysteine (Sigma-Aldrich). 10 μM Y-27632 was added for the first two days of culture. Primary organoids were cultured for 5-6 days, after which regenerative growth (number of *de novo* crypt domains per organoid) was quantified and organoids sub-cultured. Quantification was done blindly, whenever possible. Sub-culturing was performed by mechanically disrupting organoids to single crypt fragments, which were re-plated (1:4) to fresh matrigel. Secondary cultures were confirmed to start from single crypt domain by inspection, and their survival and *de novo* crypt number was quantified 2 days after re-plating. When indicated ENR media was supplemented with recombinant Aspn (rAspn) or equal amount of vehicle (PBS with 0.1% BSA) was used in controls. ENR supplemented with 10 nM Gastrin (Sigma-Aldrich), Wnt3A (RnD), 1 mM Nicotinamide (Sigma-Aldrich), and 10 μM SB202190 (Sigma-Aldrich) was used for isolated human small intestinal crypts^38^. Human small intestinal organoid starting frequency was counted at day 4. When indicated, ENR media was supplemented with 0.1 ng/ml rTgfβ1, 500 ng/ml rAspn and 1 μg/ml of ultra-LEAF™ CD44 function blocking antibody (Clone IM7; Biolegend) or equal amount of isotype control IgG2b kappa (Clone RTK4533; Biolegend).

### Decellularization

iECM was prepared by decellularizing ileum part of the mouse intestine. Briefly, intestine was flushed through with ice cold MQ water. Mucus and mesentery were removed, and incubated with MQ water overnight at +4°C. Following the incubation, intestine was flushed through with MQ water. Then, the intestine was cut into smaller pieces (~1 cm) and incubated with 1% Sodium Deoxycholate (SDC) for 3 hr at room temperature (RT) on a shaker. Pieces of the intestine were washed with MQ water for 15 min at RT on a shaker. Intestinal pieces are further incubated with 1 M Sodium Chloride (NaCl) and DNaseI (1 U/10 μl) for 2 hr at RT on a shaker. Finally, pieces of iECM were washed with PBS for 15 min at RT on a shaker before storing at +4°C (short-term) or at −80°C (long-term).

### Crypt culture on iECM

Tubular iECM was cut into open, and placed on glass bottom dish as luminal side facing upward. iECM was primed with 30 μl standard ENR for 1 hr at the standard cell culture incubator. Before plating on iECM, cultured organoids were carefully washed (3x) with cold advanced DMEM/F12. iECM was overlaid with 15-30 small round organoids (2-3 days culture) or single crypt domain (broken from passaged organoids). Standard ENR media was overlaid intermittently on iECM to avoid drying at the cell culture incubator. After 1 hr of incubation, ENR media was overlaid to the final volume of 350 μl. After overnight culture, the number of adhered organoids were counted and iECM was led to be floated off to glass using a fine tweezer. ENR media was changed every other day and crypts were counted at day 6-7 post plating for regeneration assay. When indicated, ENR media was supplemented with 500 ng/ml rAspn and 1 μg/ml of ultra-LEAF™ CD44 function blocking antibody (Clone IM7; Biolegend) or equal amount of isotype control IgG2b kappa (Clone RTK4533; Biolegend). In case of CD44 function blocking antibody or control IgG2b kappa, crypts were pre-incubated with 1 μg/ml CD44 function blocking antibody or equal amount of isotype control IgG2b kappa before plating into Matrigel, and ENR media was supplemented with 1 μg/ml of ultra-LEAF™ CD44 function blocking antibody or equal amount of isotype control IgG2b kappa in every 12 hours at the indicated duration.

### Single cell sorting

In order to isolate single cells, freshly isolated crypts were dissociated in TrypLE Express (Gibco) with 1000 U/ml of DNaseI (Roche) at +32°C for 90 seconds. Cells were washed and stained with antibodies anti-CD31-PE (Biolegend, Mec13.3), anti-CD45-PE (eBioscience, 30-F11), anti-Ter119-PE (Biolegend, Ter119), anti-EpCAM-APC (eBioscience, G8.8) and anti-CD24-Pacific Blue (Biolegend, M1/69). Cells were resuspended with SMEM media (Sigma). 10 μM 7-AAD (Life) was added to the cell suspension for live gating. Cells were sorted by using FACSAria II (BD Biosciences). Intestinal stem cells were isolated as Lgr5-EGFP^hi^; Epcam^+^; CD24^lo/−^; CD31^−^; Ter119^−^; CD45^−^; 7-AAD^−^. Lgr5^hi^ cells were cultured with standard ENR media supplemented with additional 500 μg/ml of Rspondin-1 (to yield final concentration of 1μg/ml) and 100 ng/ml Wnt3A for the first 6 days. 10 μM Y-27632 was added to the media for first 2 days. Single cell starting frequency and clonogenic growth of primary organoids were analysed at day 6-7. For flow cytometric analysis of Sca1+ cells, small intestinal epithelial cells were stained with antibodies anti-CD31-PerCP.Cy5.5 (Biolegend, Mec13.3), anti-CD45-PerCP.Cy5.5, (eBioscience, 30-F11), anti-EpCAM-BV786 (BDBiosciences, G8.8) and anti-CD24-Pacific Blue (Biolegend, M1/69), anti-Sca1-PE (Biolegend, D7), and 10 μM 7-AAD (Life). Sca1^+^ cells were isolated and analysed from the vehicle/5-FU treated young intestines (2-4 months old; 200 mg/Kg body weight) as Sca1^+^; Epcam^+^; CD31^−^; CD45^−^;7-AAD^−^.

### Real-time qPCR

RNA from crypts and cultured organoids was isolated by Trizol purification according to manufacturer’s instructions (Life). RNA from whole tissues was isolated first by homogenizing a piece of tissue in 1 ml Trizol with Precellys 24 tissue homogenizer. Isolated RNA was digested with DNasI enzyme (ThermoFisher Scientific) and transcribed with cDNA synthesis kit using OligodT primers (Molecular probes). qPCR amplification was detected by SYBRGreen (2xSYBRGreen mix, Applied biosciences) method. Samples were run as triplicates and genes of interest were normalized to *Actin/18srRNA/Rpl13a*. Primers used for qPCR-

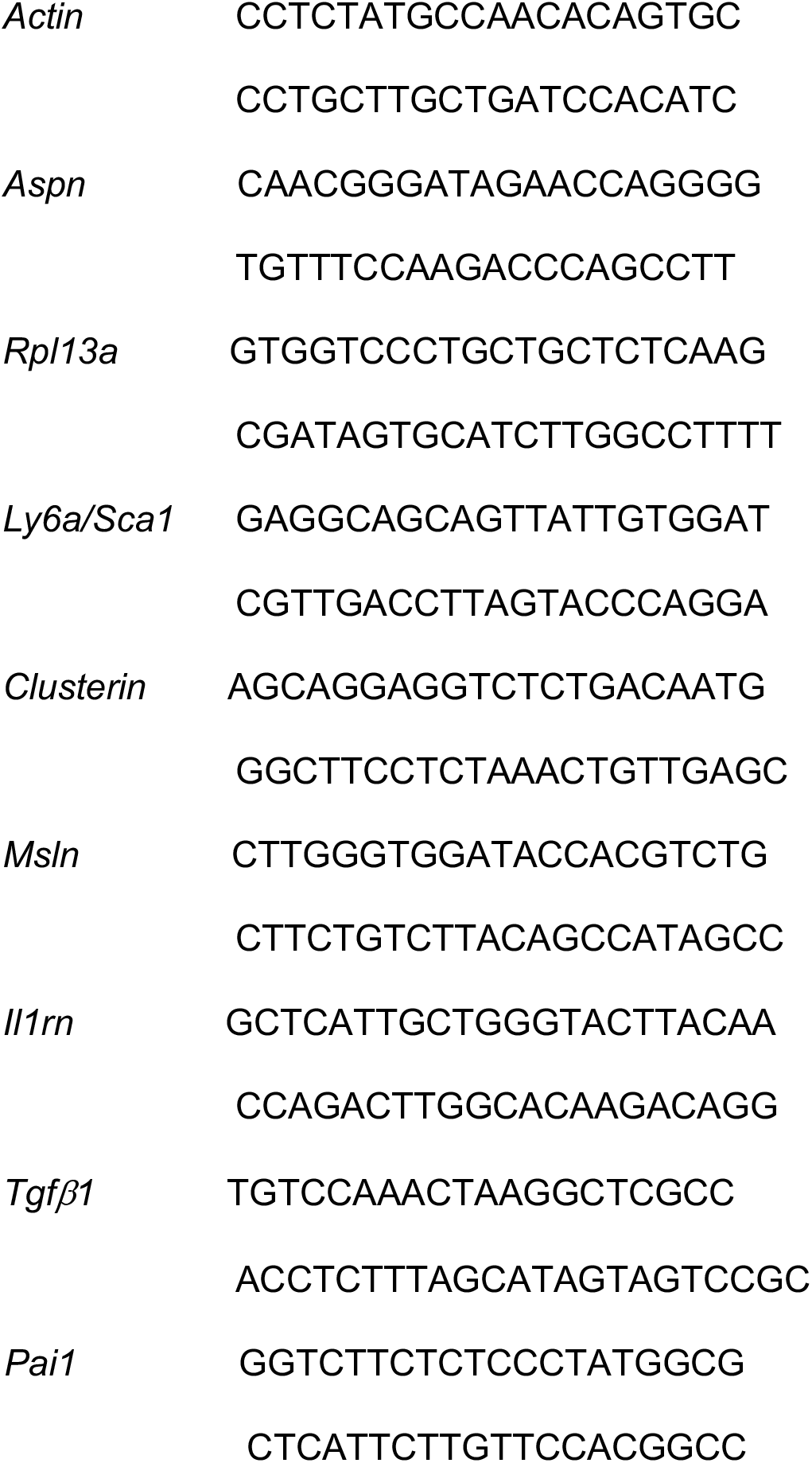

### RNA sequencing and data processing

Total RNA was isolated by using RNeasy Mini Kit (Qiagen) according to the Manufacturer’s instructions. On-Column DNase (Qiagen) digestion was performed. For RNAseq from intestinal organoids, an Ovation Universal RNA-Seq System kit was used for Illumina library preparations (NuGEN Technologies Inc., CA, USA). Purified total RNA (8.5-100 ng) was used and primers for ribosomal removal were designed and used as outlined in the kit manual. Libraries were purified with AMPure XP beads (Beckman Coulter Inc., MA, USA), quantified and run on a NextSeq 500 sequencer using 75b single read kits (Illumina, CA, USA). The read quality was examined with Fastqc 0.11.8 (https://www.bioinformatics.babraham.ac.uk/projects/fastqc/). The reads were mapped with STAR 2.5.3a^39^ to the Gencode version M16 primary genome assembly with corresponding annotation. The genome fasta and gtf files were downloaded from www.gencodegenes.org. Post-mapping quality control and gene quantification was performed with QoRTs 1.3.0^40^. R 3.6.2 (https://www.r-project.org/) was used for downstream analysis. Differential expression analysis was performed with DESeq2 1.24.0^41^, gene set enrichment analysis with camera from the package limma 3.40.2^42^. A rank-based test of enrichment was performed with camera using voom-transformed normalized counts. Hallmark and C2 gene sets from MSigDB^43^ were collected with the R package msigdbr 7.0.1 (https://cran.r-project.org/web/packages/msigdbr/index.html).

### Immunoblotting

Whole tissue samples were homogenized in RIPA buffer with 1xHalt Protease inhibitor (ThermoFisher Scientific) and 1xPhosStop (Roche) phosphatase inhibitors by using Precellys tissue homogenizer. Protein concentrations of cleared lysates were measured by DC Protein Assay kit (Bio-Rad). Samples were run on 4-12% Bis-Tris protein gels (Life) and blotted on nitrocellulose membranes. Membranes were incubated with primary antibodies: Aspn (1/1000; Sigma, SAB2500127), pSmad3 (Ser423/425) (1/1500; CST, C25A9), Smad3 (1/1500; CST, C67H9), Beta-Actin (1/5000; CST, 4967), Alpha-Tubulin (1/3000; CST, 2144) at +4°C followed by incubating with HRP conjugated anti-rabbit (Sigma-Aldrich) or anti-mouse (CST) or anti-goat (Dako) for 1 hr RT. Signal was detected using ECL reagent (ThermoFisher Scientific).

### Immunofluorescence

Tissues were fixed in 4% PFA, processed (Leica ASP200), paraffin embedded, and sectioned. Antigen retrieval was performed boiling in pH6 Citrate buffer (Sigma-Aldrich) for 20 min. Antibodies: E-cadherin (1/500; BD, 610181), pSmad2 (1/500; Abcam, ab188334), Sca1 (1/500; Biolegend, D7), CD44 (1/500; ThermoFisher, IM7). Antigen retrieval was followed by permeabilization with 0.5% Triton-X100 (Sigma). EdU incorporation was followed by EdU Click-IT chemistry according to manufacturer’s instructions (ThermoFisher Scientific). Following fixation (4% PFA), iECM was incubated with blocking buffer (5% Goat serum, 0.2% BSA, and 0.3% Triton X-100 in PBS) for 30min RT, and washed twice with PBS. Primary pan-laminin antibody (1/300; Abcam, ab11575) was diluted with blocking buffer, and incubated at +4°C overnight on a shaker at 10rpm. iECM was washed 3 times with PBS and incubated with secondary antibody for 1h at RT. After washing with PBS, samples were imaged using a spinning disc confocal. Primary antibodies were detected with biotin-conjugated secondary antibodies. For immunofluorescence Alexa-488, Alexa-594, Alexa-633 and Alexa-647 conjugated anti-rabbit or anti-mouse secondary (Life) were used. Nuclei were co-stained with DAPI (Life) or Hoechst 33342 (Life).

### Quantification of nuclear pSmad2 and EdU+ and villus cellular density

ImageJ was used for quantifying nuclear pSmad2 signal intensity. Blinded investigators measured pSmad2 mean fluorescent intensity from nuclear ROIs of cells from the nucleus of CBC and Paneth cells (identified by nuclear morphology and cellular shape) of the crypts (>20 crypts/mouse). Background subtraction was carried out based on non-nuclear pSmad2 stained area of the crypts. EdU+ cells were quantified using CaseViewer. Ileal villus cellular density (nuclei per μm) was quantified using CaseViewer.

### RNA *in situ* hybridization

RNA *in situ* hybridization was performed with RNAScope® 2.5HD Assay-Brown according to manufacturer’s protocol (RNAScope® ACDBio). Probe used: Mouse Aspn: Mm-Aspn 300031. Samples were counter stained with hematoxylin.

### Cell culture

Colorectal cancer cells HCT-116 were cultured in RPMI-1640 media (Sigma) supplemented with 10% FCS (Gibco), 100 U/ml of Penicillin/Streptomycin (Orion/Sigma) and 2 mM L-glutamine (Sigma).

### rAspn production and purification

Recombinant mouse Aspn (rAspn) was produced in Chinese Hamster Ovary (CHO) cells. Mouse Aspn expressing CHO cells were cultured in Alpha MEM (Gibco) medium supplemented with 10% Dialyzed serum (Sigma), 100 U/ml of Penicillin/Streptomycin (Orion/Sigma), 2 mM L-glutamine (Sigma), and 75 ug/ml of Zeocin (Sigma). For harvesting rAspn containing media supernatant, confluent cells (>80%) were cultured without Zeocin for 3 days in alpha MEM medium supplemented with 2 mM Glutamine, 100 U/ml of Penicillin/Streptomycin (Orion/Sigma) and 1% FCS (Gibco). Followed by 3 days of culture, media supernatant was collected, centrifuged, and stored at −80°C. Finally, rAspn was purified from the supernatant using standard purification procedure. 500 ng/ml rAspn was used in culturing intestinal epithelium.

### Mass spectrometry

Proteomic samples from old iECM are prepared in 6M Urea and analyzed similarly as published ^44^ before. MaxQuant (1.6.10.43) database search was used for peptide and corresponding protein identifications (Data S2).

### Statistical analysis

For analysis of *in vitro* organoid culture, crypt culture on iECM, and histological quantification investigators were blinded when possible. Microsoft Excel and Graphpad Prism were used for statistical analysis and visualization of data. All data were analysed by two-tailed Student’s t-test. Paired t-test was applied when appropriate and noted in the figure legends. Statistical significance of the overlap between two groups of genes for Fig. 3b was calculated using the online tool (http://nemates.org/MA/progs/overlap_stats.html).

### Human Biopsy samples

Human jejunal samples were obtained from patients undergoing Roux en-Y gastric bypass surgery. The tissue samples used for organoid functional assay were stored in normal saline on ice until crypt isolation. The study regarding relevant samples was approved by Helsinki University Hospital. Written and informed consent was obtained prior to enrolment.

### Animals

*Lgr5-EGFP-IRES-CreERT2^45^, Twist2-Cre;Aspn^fl/fl^, Twist2-Cre;R26R^LSL-tdtomato/+^* mice were kept in C57BL/6 background. For *in vivo* proliferation assessment, EdU (20 mg/Kg) in PBS was injected intraperitoneally 2 hours prior to sacrificing the mice. 5-Fluorouracil (Sigma) was reconstituted in DMSO (100 mg/ml) and single intraperitoneal injection was given to the mice with a dose of 200 mg/Kg body weight. Young mice with 3-6 months of age were considered young, and used for all the experiments. Mice with targeted reporter allele for *Aspn* (*Aspn^Lacz^*; C57BL/6N-Aspntm1a(EUCOMM)Hmgu/Ieg) were purchased (INFRAFRONTIER/EMMA), and reporter allele was converted into conditional allele (*Aspn^lox^*) by crossing with mice expressing FLP recombinase under CAG promoter. Genotyping of the mice were carried out with described primers (Table-2). Animal housing and all the animal experiments were approved and carried out in accordance with the regulations of Finnish national animal experimentation board.

### Electron Microscopy

iECM was fixed with 2% Glutaraldehyde in 100 mM Na-Cacodylate (NaCac) buffer (pH 7.4) for 1h at room temperature. Samples were then osmicated with 1% O_s_O_4_ in 0.1M NaCac followed by several washings with 0.1M NaCac and dH_2_O before the samples were dehydrated and dried overnight. Samples are platinum coated and scanning electron micrographs were obtained using FEI Quanta 250 Field Emission Gun SEM.

## Data and materials availability

sequencing and tissue mass spectrometry data is publicly available through ArrayExpress (upon publication).

## Acknowledgments

We thank Professor Marja Mikkola and Professor Ari Ristimäki for their guidance in thesis committee for S.I., and J. Bärlund and M. Simula for technical supports. Light Microscopy and Electron Microscopy Units, Meilahti Clinical Proteomics Core Facility, DNA Sequencing and Genomics Unit of University of Helsinki, and Laboratory Animal Center, University of Helsinki are thanked for their services.

## Author contributions

S.I. and P.K. designed and interpreted the results of all the experiments. S.I., E.N., S.A., N.P., A.K., A.W., M.V. performed all the experiments and analysed the results. D.B. processed and analysed the RNA-sequencing data. E.N. and S.I. analysed the immunofluorescence images. T.S., A.J. and K.H.P. provided the human biopsy material. A.O., K.B.J and M.O. participated in the design and interpretation of experiments. S.I. and P.K. wrote the paper.

---

**Extended Data Figure 1.**
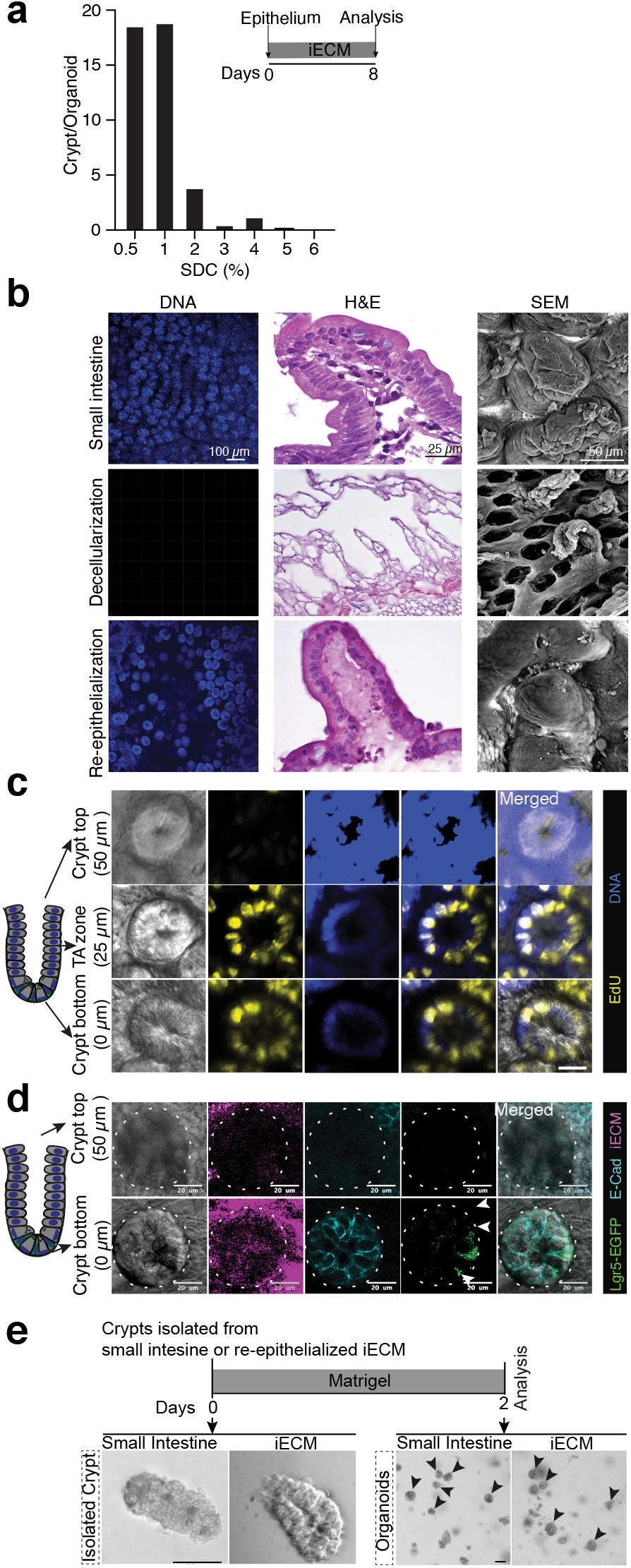
Organotypic growth of intestinal epithelium on iECM. **a**, Optimization of the amount of sodium deoxycholate (SDC) for iECM generation. Different amount of SDC (0.5-6%) was used for the tissue decellularization. Number of crypts/organoid was quantified 8 days after seeding epithelium on iECM. **b**, Decellularization of mouse small intestine preserves the ECM contour of crypts and villi. Repopulation of crypts and villi guided by the iECM. DNA: Hoechst stain, H&E: Haematoxylin+Eosin stain of cryosections, SEM: scanning electron micrograph. **c**, Proliferation on iECM is restricted to the repopulated crypt pits. Representative images of Edu+ proliferative cells at the crypt base. Confocal fluorescence microscopy. Blue (DNA, Hoechst), Yellow (EdU). Scale bar 20 μm. **d**, Lgr5-EGFP cells are restricted to the crypt base on iECM. Representative fluorescent images of iECM repopulated with epithelium from *Lgr5-CreERt2-IRES-EGFP* mouse. Pink (iECM, autofluorescence of collagen), Cyan (E-Cad), Green (EGFP labeled Lgr5 expressing stem cells). Scale bar 20 μm. All the images were obtained after 7 days of culture on iECM. **e**,Comparison of morphologies of the freshly isolated crypts (left) and matrigel-cultured organoids (right; 2 days in culture), obtained from small intestine and re-epithelialized iECM (7 days culture). Scale bar 50 μm.

**Extended Data Figure 2.**
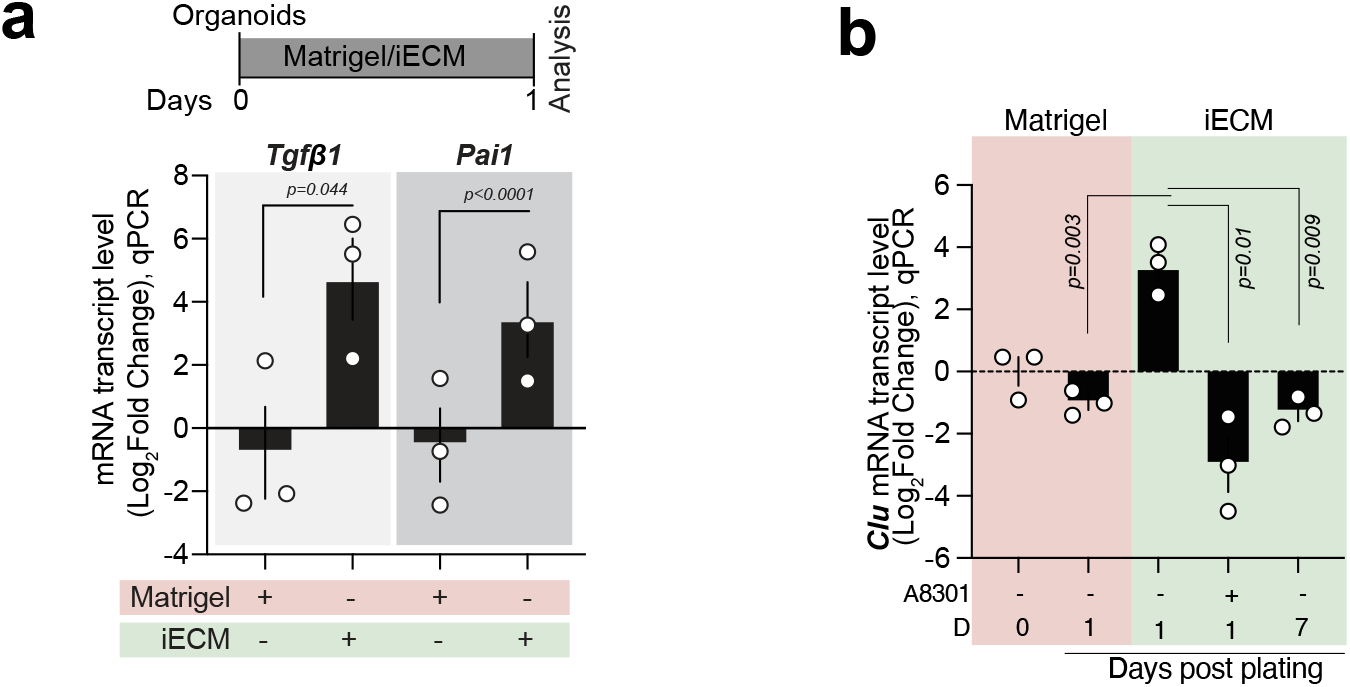
Comparison of the induction of the Tgfβ pathway-responsive genes, and marker of revival stem cells, in matrigel and iECM. **a**, qPCR analysis of *Tgfβ1 and Pai1* expression in the intestinal epithelium after 24 hours culture in matrigel and on iECM. Values show fold change in comparison to matrigel-cultured organoids. *Rpl13a* was used as a reference gene. Student’s paired *t*-test, mean +/− s.e.m. **b**, qPCR analysis of the relative mRNA level of *Sca1* from the intestinal organoids (D0: before passaging; D1: 24 hrs after passaging in matrigel) and intestinal epithelium on iECM (D1 & D7: Day 1 & Day 7 post-plating on iECM from matrigel). *Rpl13a* was used as a reference gene. Values show fold change in comparison to Day 0 intestinal organoids. Student’s paired *t*-test, mean +/− s.e.m.

**Extended Data Figure 3.**
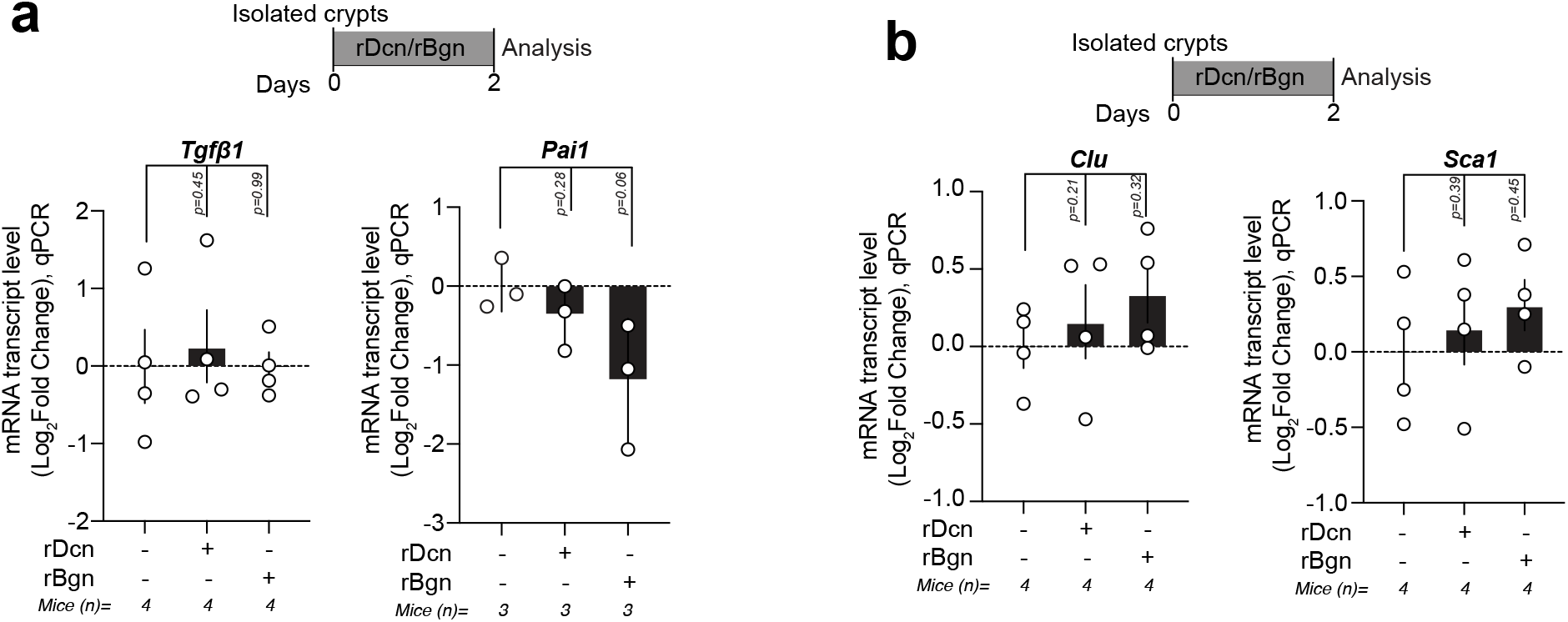
Effects of the recombinant Decorin (rDcn) and Biglycan (rBgn) in the expression of the marker genes of the Tgfβ pathway and fetal state in the intestinal organoids. (**a**-**b**) qPCR analysis of *Tgfβ1 and Pai1* expression (in **a**) – and *Clu* and *Sca1* expression (in **b**) in the rDcn and rBgn-treated (48 hrs) intestinal organoids. Values show fold change in comparison to untreated control organoids. *Rpl13a* was used as a reference gene. Student’s paired *t*-test, mean +/− s.e.m.

**Extended Data Figure 4.**
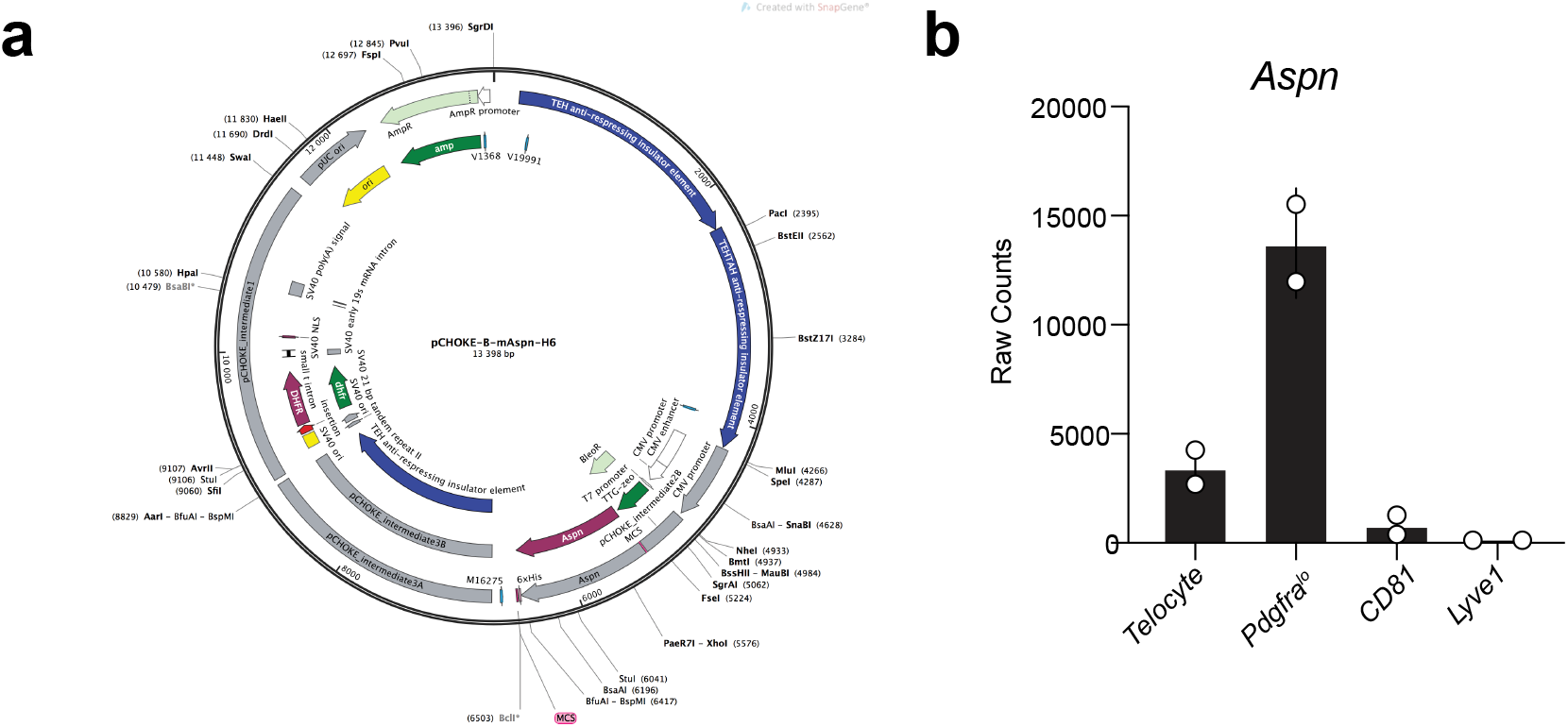
Production of recombinant Aspn (rAspn), and *Aspn* expression in the intestinal mesenchyme. **a**, Construct used to produce recombinant Aspn (rAspn) in CHO cells. **b**, Expression of *Aspn* in the different mesenchymal cell types of mouse small intestine. Data was obtained from the previously published publically available database (GEO accession number: GSE130681).

**Extended Data Figure 5.**
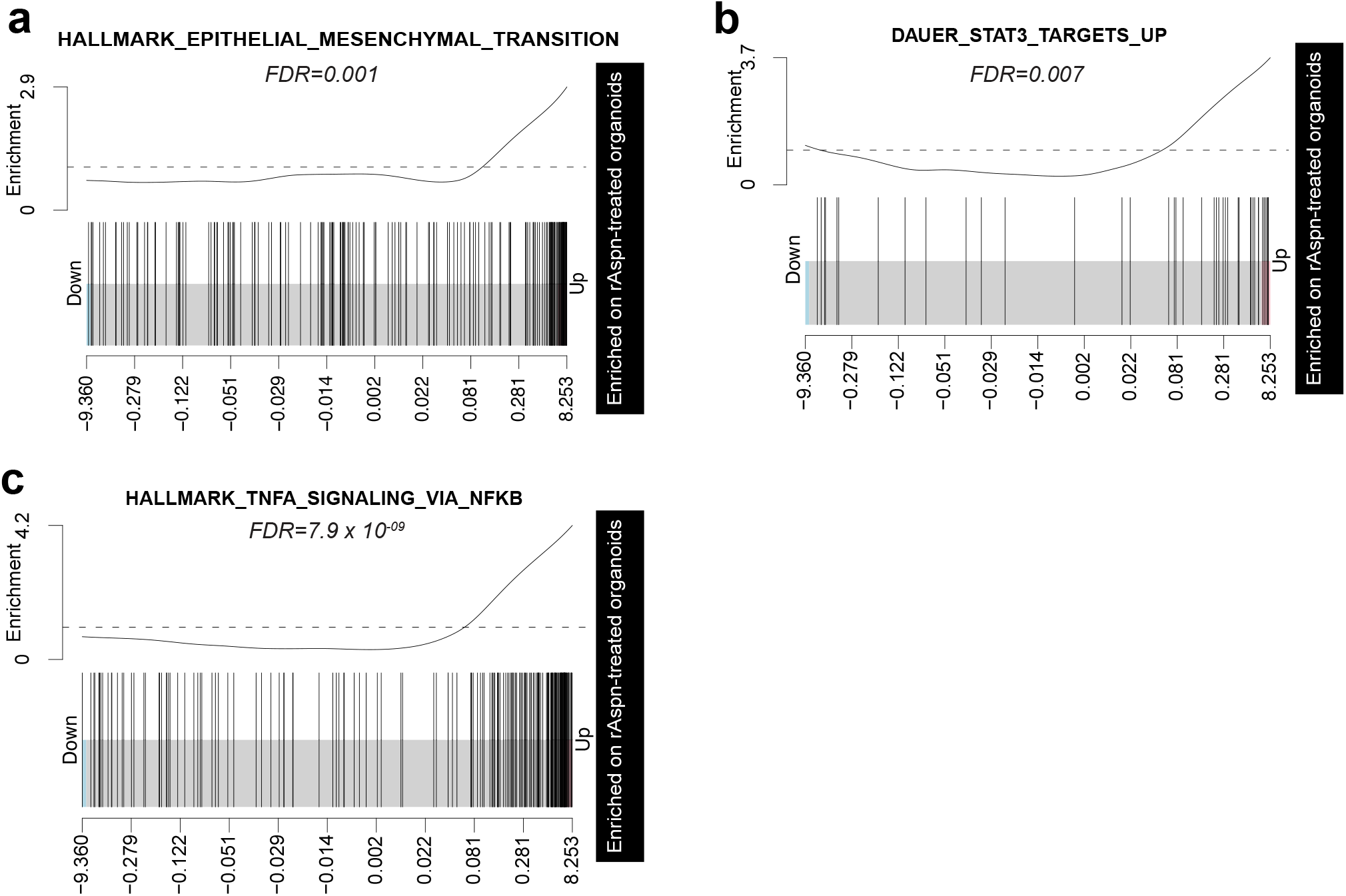
Enrichment of CD44 downstream signalling pathways in rAspn-treated organoids. **a-c**, GSEA analysis of the gene list “Hallmark_Epithelial_Mesenchymal_Transition (**a**), Dauer_Stat3_Targets_Up (**b**) and Hallmark_Tnfa_Signalling_via_Nfkb (**c**)” in the intestinal organoids treated with rAspn (500 ng/ml; 48 hours). False discovery rate (FDR) is shown.

**Extended Data Figure 6.**
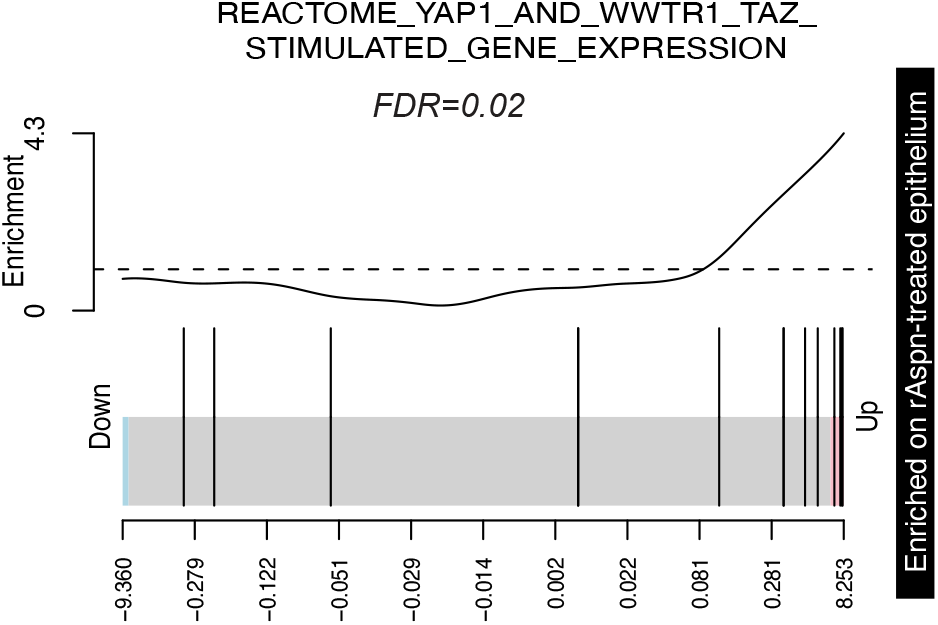
Yap1/Taz pathway-related gene expression in the rAspn-treated intestinal organoids. GSEA analysis of the gene list “REACTOME_YAP1_AND_WWTR1_TAZ_STIMULATED_GENE EXPRESSION” in the intestinal organoids treated with rAspn. False discovery rate (FDR) is shown.

**Extended Data Figure 7.**
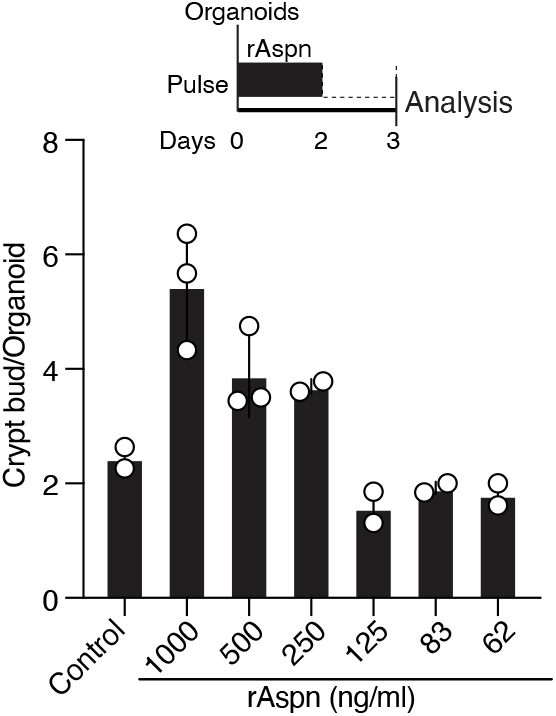
Titration of the doses of rAspn for treating intestinal organoids. Titration of rAspn concentrations for effects on intestinal organoids (n=2-3 mice). Untreated samples were used as control. Mean +/− s.d.

**Extended Data Figure 8.**
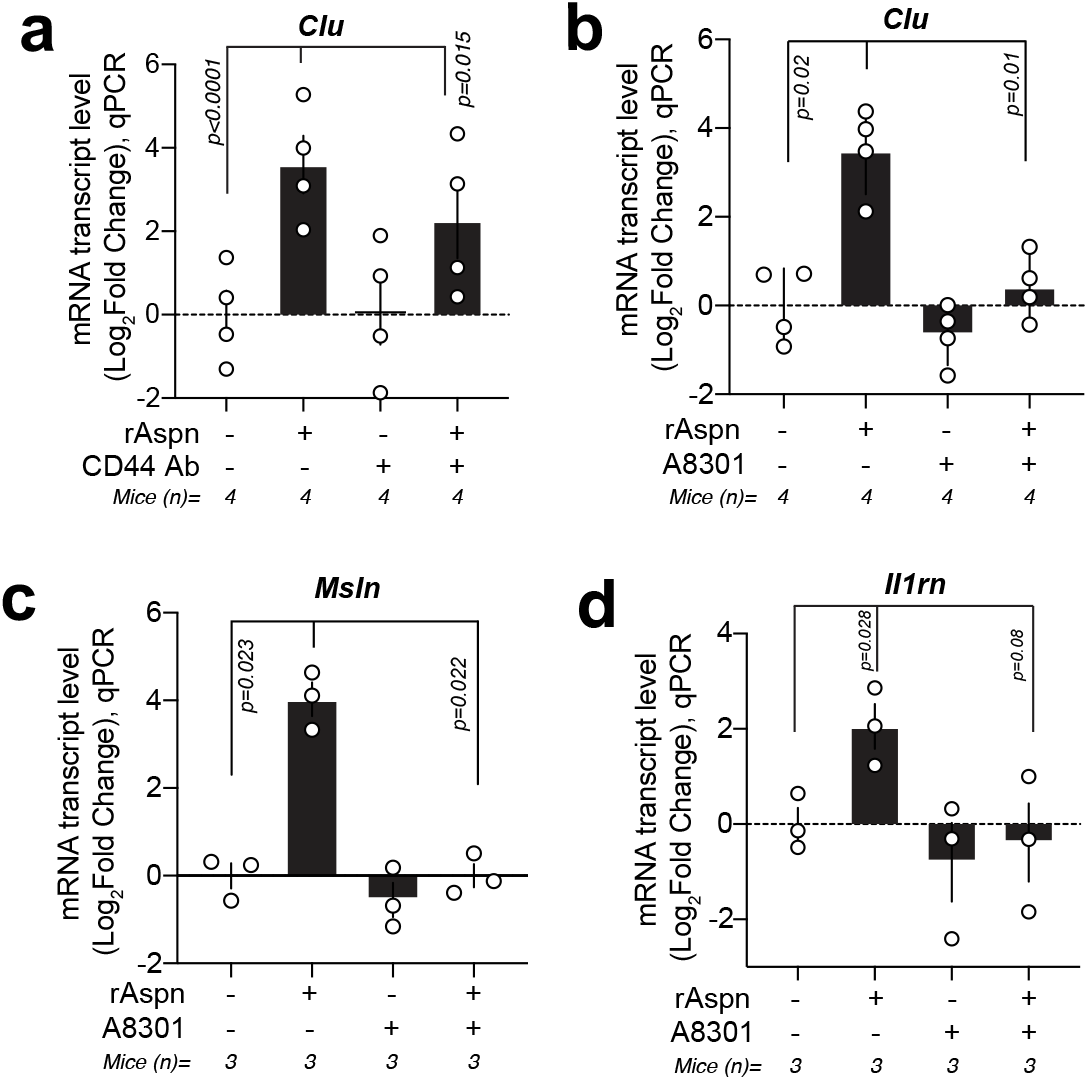
rAspn induces fetal-like transcription TgfbRI dependently. **a**, qPCR analysis of *Clusterin* (*Clu*) in rAspn and/or CD44 function-blocking antibody treated small intestinal orgnaoids (n=4). Isotype control IgG2b kappa was used in samples not receiving CD44 Ab. Values show fold change in comparison to untreated control organoids. *Rpl13a* was used as a reference gene. Mean +/− s.e.m. Student’s paired *t*-test. **b-d**, qPCR analysis of the relative mRNA level of *Clu, Il1rn*, and *Msln*, in the rAspn and/or TgfβType I receptor inhibitor (A8301)-treated mouse intestinal organoids (n=4 mice for Clu, n=3 mice for *Msln*, and *Il1rn*). Values show fold change in comparison to untreated control organoids. *Rpl13a* was used as a reference gene. Mean +/− s.e.m. Student’s paired *t*-test.

**Extended Data Figure 9.**
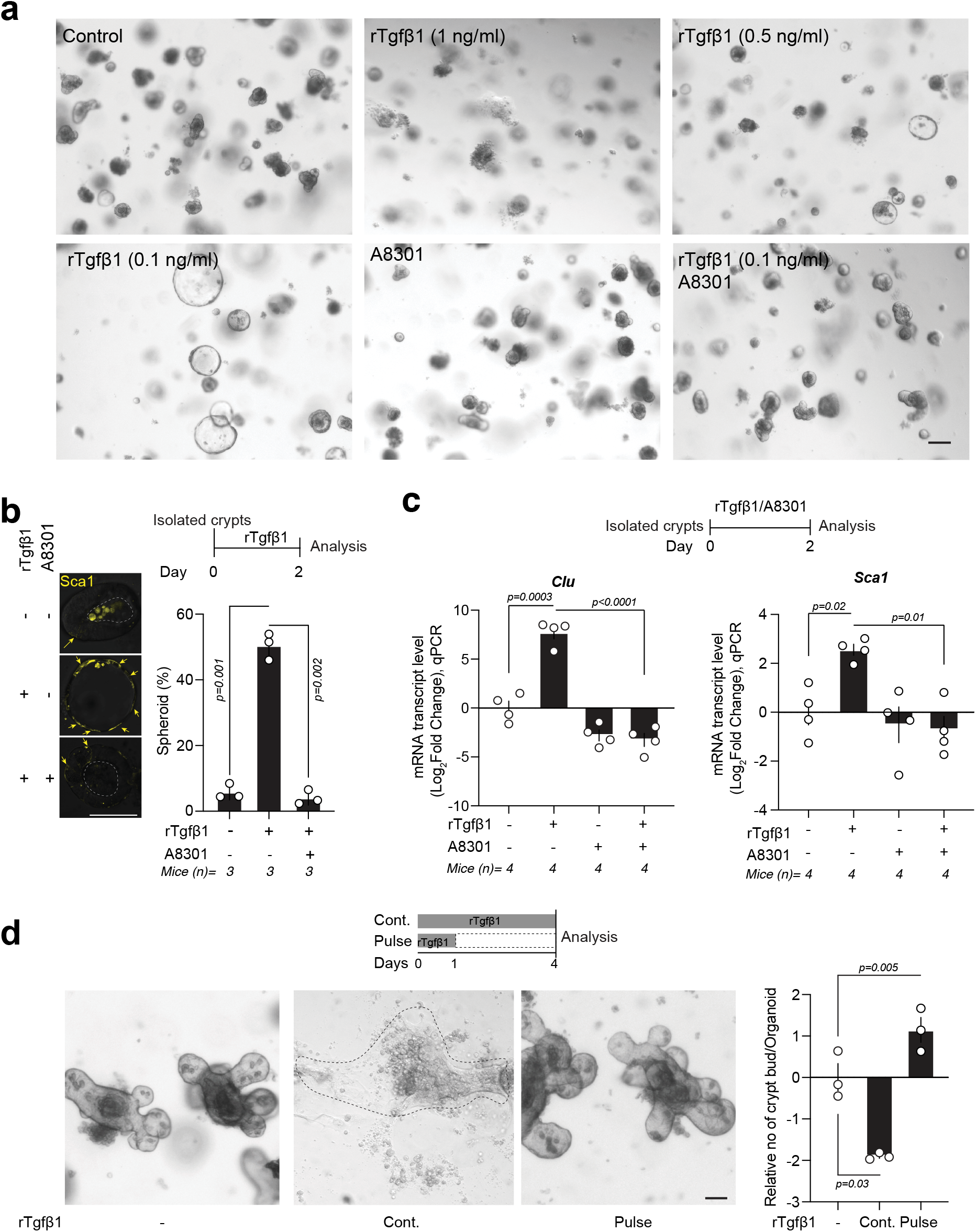
rTgfβ1 induces fetal-like state in the intestinal organoids. **a**, Representative micrographs of the intestinal organoids with different concentrations of the rTgfβ1 with/without Tgfβ type I receptor inhibitor A8301 (500 nM). Isolated crypts were cultured under the indicated conditions in the micrographs for 2 days. Scale bar 50 μm. **b**, Analysis of spheroid forming capacity of rTgfβ1 (0.1 ng/ml) and/or A8301 (500 nM) treated mouse intestinal organoids (left; n=3). Mean +/− s.d. Student’s paired *t*-test. Scale bar 20 μm. **c**, qPCR analysis of *Clu* and *Sca1* from rTgfβ1 (0.1 ng/ml) and/or A8301 (500 nM) -treated intestinal organoids (n=4). Values show fold change in comparison to untreated control organoids. *Rpl13a* was used as a reference gene. Mean +/− s.e.m. **d**, Regenerative growth of crypts (n=3 mice) with transient (1 day) and sustained (5 days) treatment of rTgf β1 (0.1 ng/ml).Values in the bar graph were normalized with the untreated control samples. Mean +/− s.e.m. Scale bar 50 μm. Student’s paired *t*-test.

**Extended Data Figure 10.**
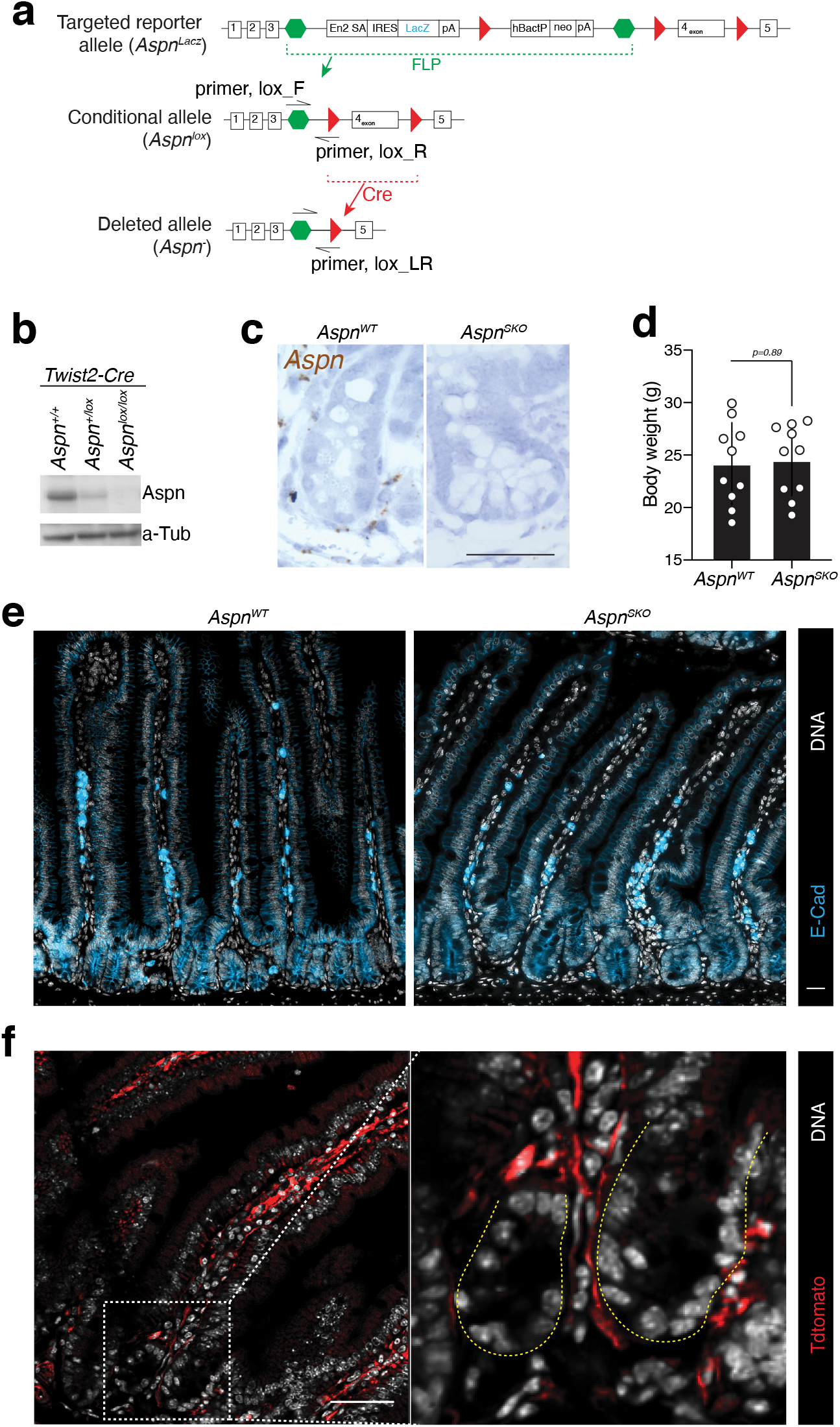
Generation of a mouse model with conditional *Aspn* allele, and tissue specific deletion of mesenchymal *Aspn*. **a**, Targeting and development of *Aspn* alleles. **b**, Immunoblot of Aspn and alpha-Tubulin of the ileal tissue samples from the *Twist2-Cre; Aspn^+/+^ (Aspn^WT^)*, *Twist2-Cre; Aspn^+/lox^* & *Twist2-Cre; Aspn^lox/lox^ (Aspn^SKO^)* mice. **c**, *In situ* analysis of *Aspn* expression in *Aspn^WT^* and *Aspn^SKO^* mice. Scale bar 50 μm. **d**, Body weight analysis of the young (2-6 mo) *Aspn^WT^* and *Aspn^SKO^* littermates (n=10 pairs mice). Loss of mesechymal *Aspn* has no adverse effects on growth. Mean +/− s.d. Student’s unpaired *t*-test. **e**, Comparison of the small intestinal morphology from *Aspn^WT^* and *Aspn^SKO^* mice. White (DNA), cyan (E-Cad). Scale bar 50 μm. **f**, Lineage tracing in the *Twist2-Cre; R26R^LSL-tdtomato/+^* mice. Red cells show the Cre-mediated recombined cells. Yellow dotted lines mark the crypt epithelium. White (DNA), Red (Tdtomato). Scale bar 50 μm.

**Extended Data Figure 11.**
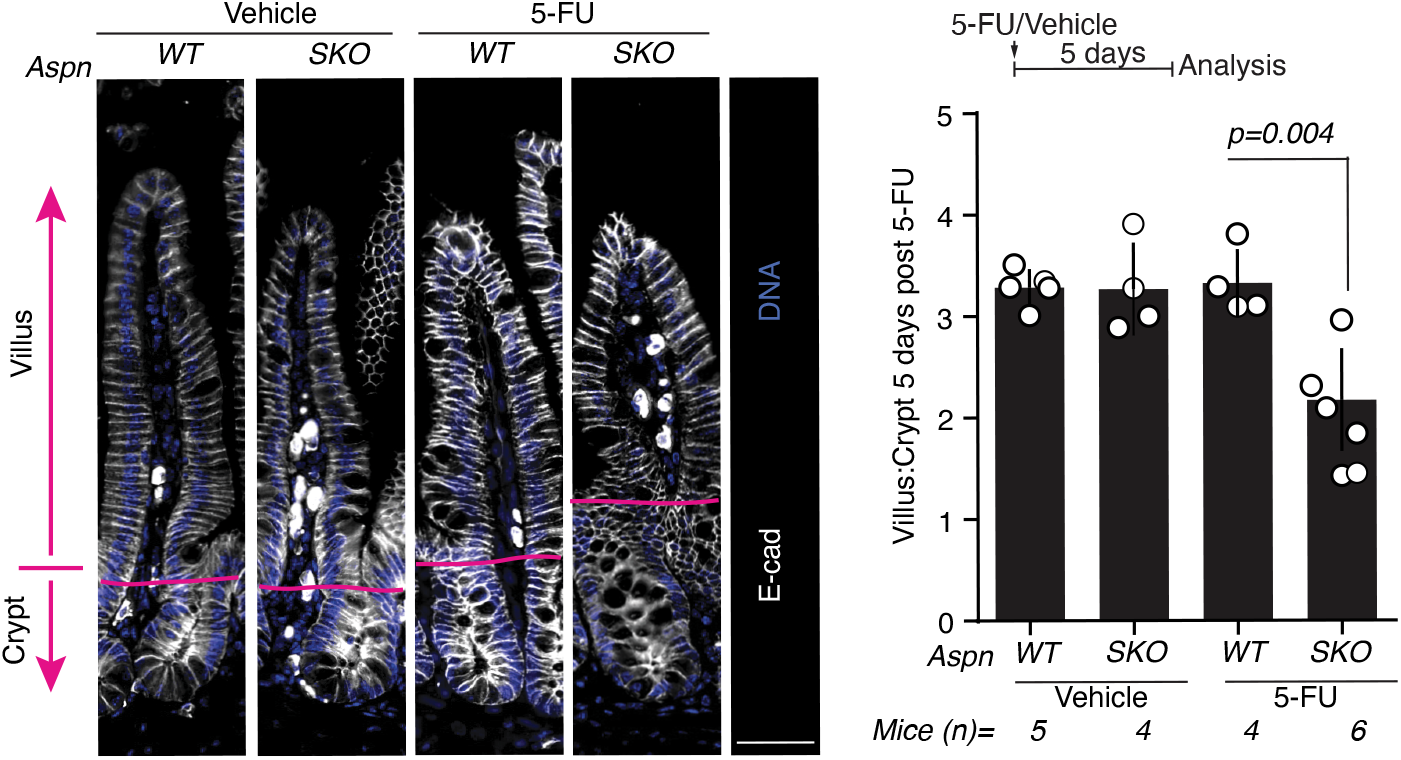
Conditional knock out of *Aspn* impedes intestinal regeneration. Analysis of crypt and villus lengths in the histological sections of *Aspn^WT^* and *Aspn^SKO^* mice five days after treatment with 5-FU (200 mg/kg body weight) or vehicle only (DMSO). White (E-Cad), Blue (DNA). Mean+/−s.d. Scale bar 50 μm. Student’s unpaired *t*-test.

**Extended Data Figure 12.**
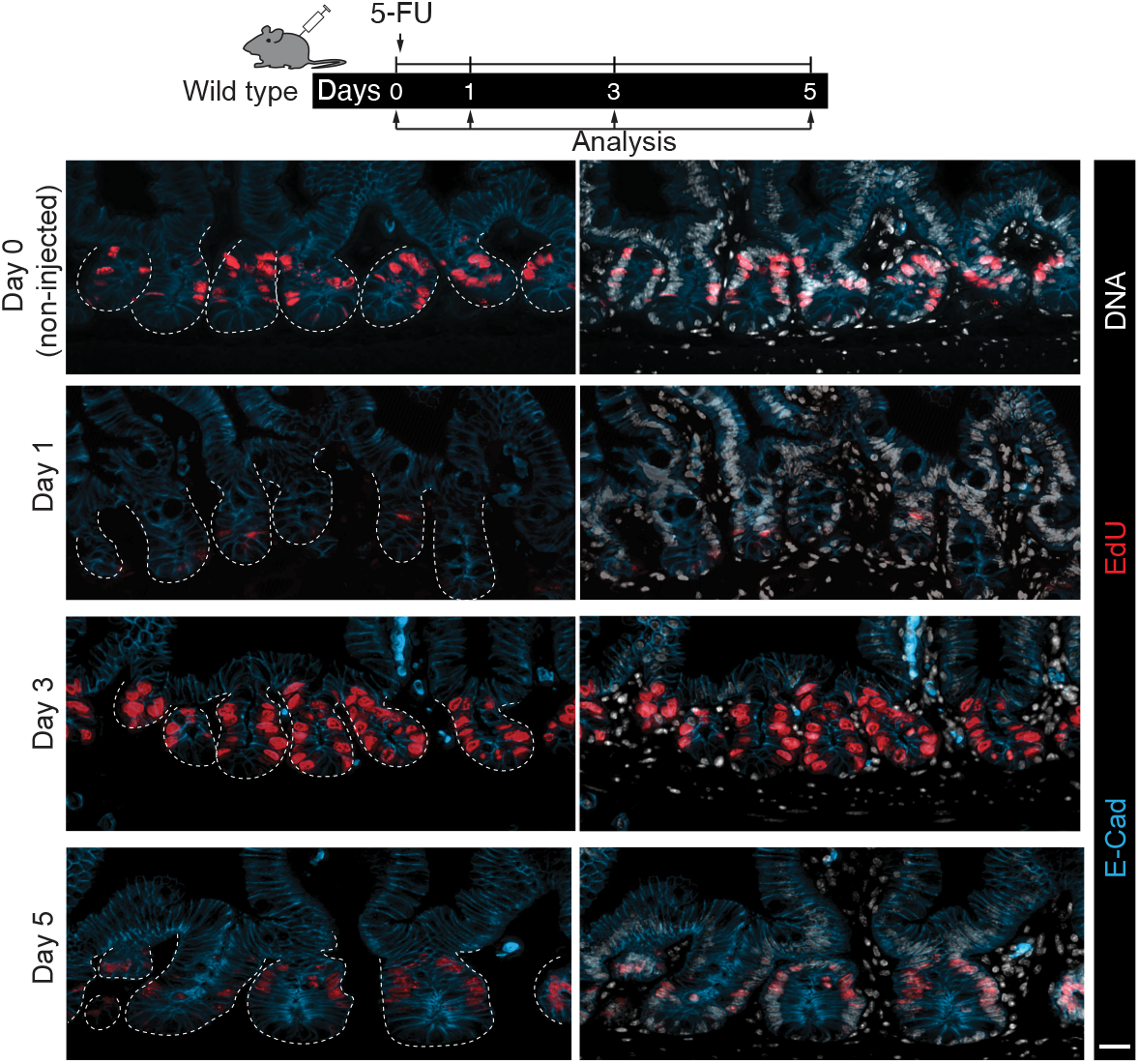
Temporal dynamics of the proliferative cells in the 5-FU induced regenerating intestine. Comparison of the intestinal epithelial proliferation upon 5-FU injection (200 mg/Kg body weight) at different time points - Day 0 (non-injected), Day 1, Day 3 and Day 5 post injections in wild type mice. White (DNA), Red (EdU), Cyan (E-Cad). Scale bar 20 μm.

**Extended Data Figure 13.**
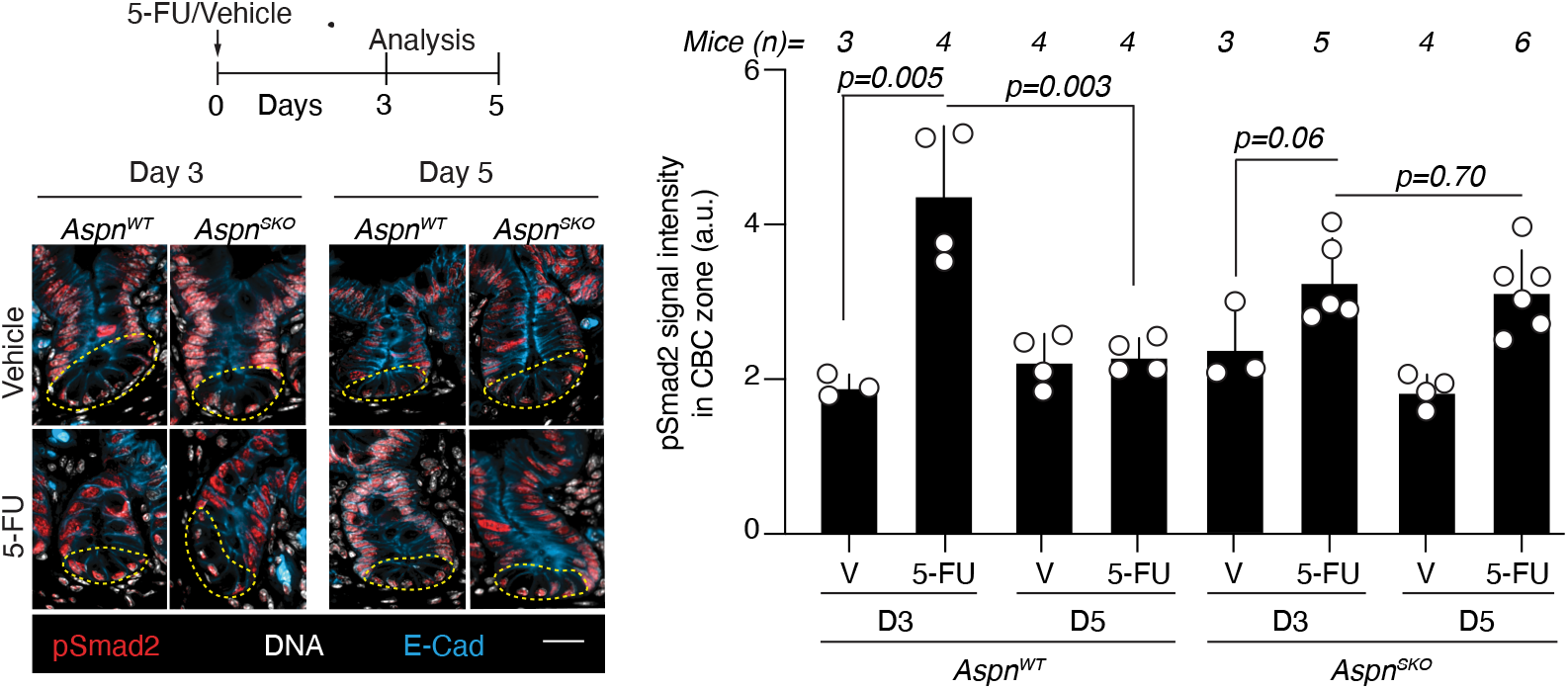
Induction dynamic of epithelial Tgfb signaling is altered in mice with mesenchymal deletion of Aspn. pSmad2 staining (Red) of the crypt base cells three (D3) and five (D5) days after Vehicle (V) or 5-FU (200 mg/kg body weight) injection in *Aspn^WT^* and *Aspn^SKO^* mice. (n=3-6 mice per group). White (E-Cad), Blue (DNA). Mean+/−s.d. Scale bar 50 μm. Student’s unpaired *t*-test.

**Extended Data Figure 14.**
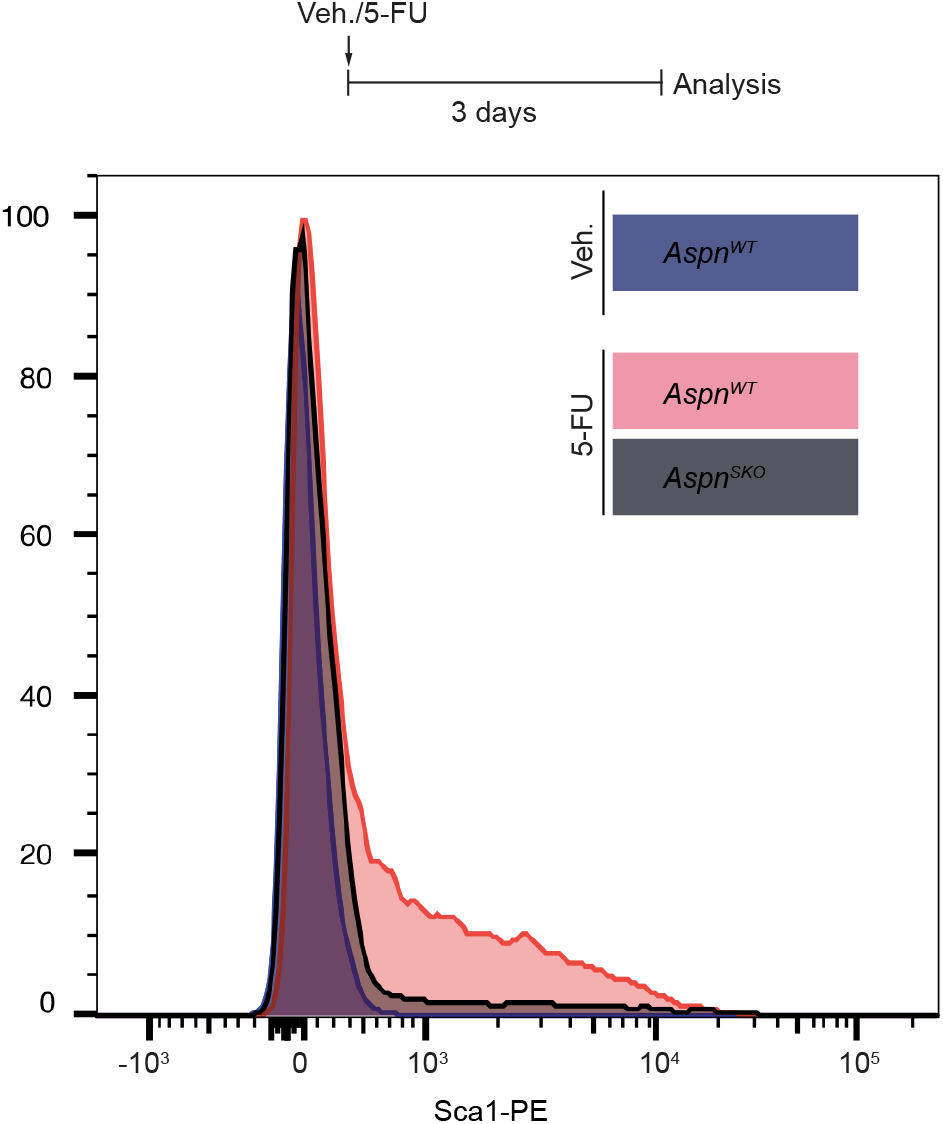
Damage-induced increase in Sca1+ cells is dimished in *Aspn^SKO^* mice. Flow cytometric analysis of small intestinal Sca1^+^ EpCam^+^ CD31^−^CD45^−^ cells isolated from vehicle and 5-FU treated *Aspn WT* and *Aspn^SKO^* mice (n=2-5; Day 3 post injection). Histogram shows distribution of Sca1 intensity in representative examples.

